# Engineering elephant models of cold adaptation and cancer resistance

**DOI:** 10.1101/2024.09.07.611789

**Authors:** Emil Karpinski, Nikil Badey, Esther Mintzer, Asaf Ashkenazy-Titelman, Li Li, George M. Church

**Affiliations:** Department of Genetics, Harvard Medical School, Boston, MA, 02115, USA; Wyss Institute for Biologically Inspired Engineering, Harvard University, Boston, MA 02215, USA

**Keywords:** woolly mammoths, Asian elephants, arctic adaptation, TP53, retrogenes, CRISPR-Cas9, RNA-seq, gene regulation

## Abstract

Proboscideans have developed a suite of genetic changes including those responsible for tumor suppression and environmental adaptation. Here we examine a handful of woolly mammoth-specific deletions, and their potential contributions to arctic adaptation, as well as the expanded TP53 genetic repertoire in elephants. We use CRISPR-Cas9 to introduce mammoth-specific noncoding deletions as well as knockouts of TP53, all 29 TP53 retrogenes, or both in combination in Asian elephant cell lines, and examine the transcriptomic response. We find that many of the mammoth-specific deletions likely contribute to various arctic phenotypes including vascular development, metabolism and thermogenesis, and hair and skin adaptations. We also find that while there is considerable overlap in DNA damage responses of the TP53 and retrogene knockouts, retrogenes knockouts also exhibit strong enrichment of many extracellular pathways suggesting they may play a role in the tumor microenvironment and mitigating metastatic growth.

## Introduction

Gene regulation plays a significant role in the ability of species to adapt to novel environments and is believed to contribute to speciation via hybrid dysfunction (*1*, *2*). Adaptive regulatory changes have been identified across Animalia, including in primates, mice, birds, fish, and cervids (*2–5*). One of reasons for the large impact of regulatory mutations is their ability to fine-tune expression of genes during different developmental stages or tissues, in comparison to the near ubiquitous change that mutating protein sequences would have (*5*). However, these tissue-specific effects, as well as the fact that regulatory sequences may be very distant from their regulatory targets, make them difficult to examine bioinformatically, and necessitate the use of cell or animal models (*2*).

Throughout their ∼60-million-year evolutionary history, proboscidean evolution has largely followed two main trends: a general increase in size from the earliest proboscideans, *Eritherium azzouzorum* and *Phosphatherium escuillei* (∼5 and 17 kg respectively); and adaptive radiation to expand into and occupy many different environments from tropical wetlands to the cold tundra (*6*, *7*) (Fig. 1A). These trends are especially apparent in the elephants (*Elephantidae*), which during the late Pleistocene included one of the largest land mammals of all time, *Palaeoloxodon antiquus* (>13 tons), and species found in various environments across Africa, Eurasia, and North America (*6*, *7*).

**Figure 1.**
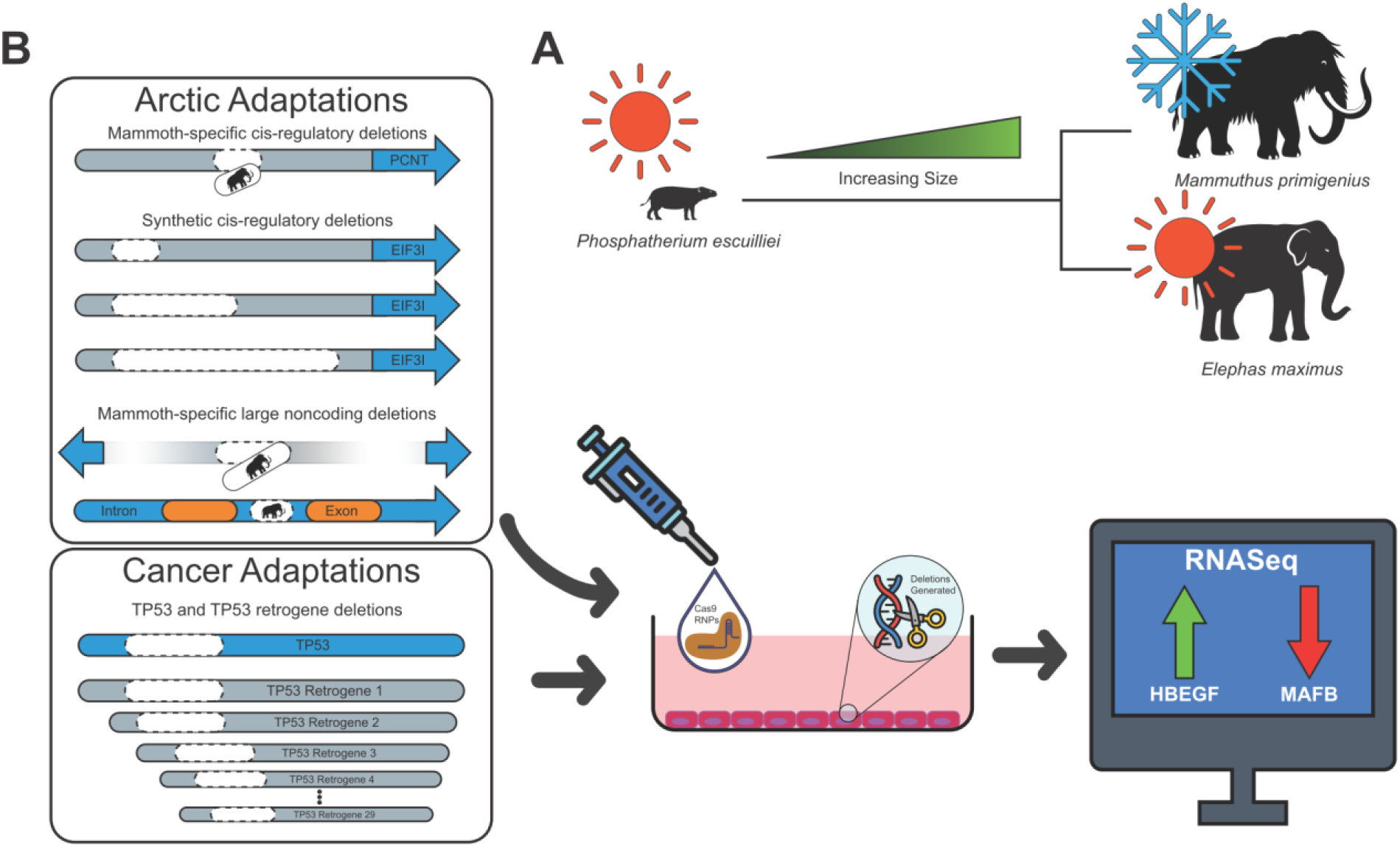
Experimental Design. **(A)** Trends in proboscidean evolution include a general increase in body size, and adaptations to various environments including warm tropical grasslands and cold arctic tundra. **(B)** We engineered mammoth specific or synthetic cis-regulatory deletions, large mammoth-specific deletions in both intra and intergenic regions, as well as deletions within TP53 and/or all TP53 RTGs using Cas9 RNPs within various Asian elephant cells in culture, before examining their effects via RNA-seq.

Woolly mammoths (*Mammuthus primigenius*) occupied dry, cold steppe-tundra environments during the coldest periods of the Pleistocene (*6*, *8*). They evolved many physiological adaptations to survive in this environment, including a multi-layered and thicker hair coat, changes in skin and fat distribution, smaller ears and tails to minimize heat exchange, and a general increase in size and growth rate. These changes are especially striking given their closest living relative, the Asian elephant (*Elephas maximus*), is primarily found in warm topical forests and grasslands environments throughout Southeast Asia (*6*, *8*). Previous studies have reported protein-coding changes between woolly mammoths and extant elephants which may have contributed to some of these phenotypes (*9*, *10*). However, the evolved regulatory landscape of woolly mammoths has been largely ignored.

Peto’s paradox is the observation that cancer incidence in a species does not always scale with body mass and/or lifespan (*11*, *12*). Elephants have emerged as a key species in attempts to resolve this paradox due to their large size, long lifespan, and very low cancer risk (*12*, *13*). Genomic analyses have identified the duplication of many tumor suppressor genes across extant elephants, including multiple retrogene copies of TP53 (*12*, *13*). Although there is still considerable uncertainty about how or which of these TP53 retrogenes (RTGs) may confer tumor resistance, previous works have shown that the encoded protein of at least one of these retrogenes, African elephant RTG 9, induces strong cellular responses to DNA damage (*12–14*). However, most previous studies have largely focused only on African elephant TP53 retrogenes and often in non-native genetic (e.g. plasmid expression) or cellular backgrounds (e.g. human or mouse cells).

In this study, we examine the regulatory changes responsible for adaptation and tumor resistance in woolly mammoths and Asian elephants in vitro. We have engineered a handful of mammoth-specific non-coding mutations into Asian elephant cells, and observed changes implicated in cardiac development, hair and skin function, and lipid metabolism. We further validate our cell culture models by examining the best-studied elephant genetic pathway – their expanded TP53 repertoire. We used CRISPR-Cas9 to disrupt TP53, all TP53 retrogenes, or both in combination in Asian elephant fibroblasts, and examined the transcriptional profiles of these knockouts with and without DNA damage stress, to resolve the protective roles of TP53 retrogenes (Fig. 1). Our results suggest that these cell culture models are tremendously useful for investigating the potential effects of both adaptive regulatory changes and tumor suppressor duplications, and generate a strong foundation of testable hypotheses for future research.

## Results

### Identification of mammoth deletions

Our in silico TFBS annotation predicted ∼1.98 million transcription factor binding sites (TFBS) upstream of 21,808 genes (mean 90.6/gene). We originally identified ∼235k mammoth-specific deletions, with an average size of 236 bp (min: 25; max: 13.2kb), of which 1,798 intersected our predicted transcription factor binding sites (TFBS; Supplementary Table 3). Using previously established methods we also predicted 629 large mammoth-specific deletions (i.e. >=500 bp; Supplementary Table 4) of which 46 were predicted to have a genome-unique guide within 25bp of their start and end coordinates (*15*). Notably we find strong concordance between both methods, with deletions identified using our own methodology intersecting 88.1% of the large mammoth deletions identified using the established pipeline.

### Mammoth-specific and Synthetic Regulatory Editing

We attempted to introduce one mammoth specific deletion upstream of PCNT (pericentrin), and one synthetic deletion upstream of EIF3I (eukaryotic translation initiation factor 3 subunit I) due to its high expression in our mesenchymal stem cell (MSC) line. PCNT mutations have implicated in the development of smaller cranial features and olfactory development (*8*, *16*), which may contribute to arctic adaptations present in woolly mammoths. While we were successful in generating mutations in these regions (Fig. 2A), we didn’t recover differential expression of PCNT in either our pilot or verification experiments. We did however observe differential expression of EIF3I, but only with 50% of the guide combinations, one of which we successfully verified. Our lack of signal in PCNT and sometimes in EIF3I underscores the effect tissue or timing has on gene regulation, and may be why we fail to observe a signal in all cases.

**Figure 2.**
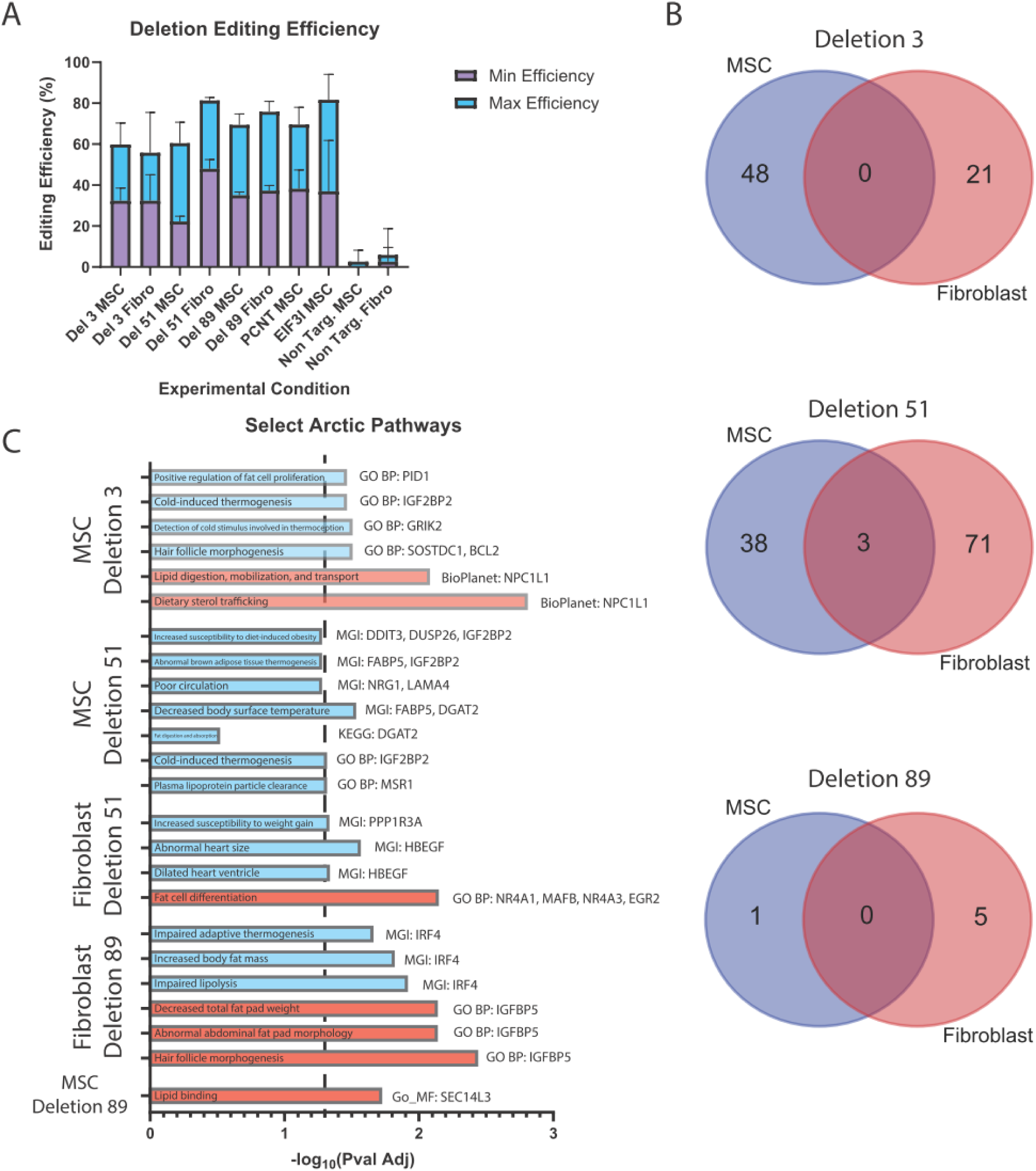
Mammoth deletion efficiency and effects. **(A)** Editing efficiency observed for each deletion in both MSCs and fibroblasts estimated using gel electrophoresis. **(B)** DEG overlap between MSCs and fibroblasts for each of the three large mammoth-specific noncoding deletions. **(C)** A selection of pathways potentially related to arctic adaptation. Bars are colored blue/red for upregulated and downregulated genes respectively. The database and key genes are listed beside each bar. A vertical dashed line indicates a <0.05 significance level.

### Mammoth-specific deletion differentially expressed genes

We also generated three large mammoth deletions (Deletions 3, 51, and 89) with comparable editing efficiency in both fibroblasts and mesenchymal stem cells (Fig. 2A). Following bulk RNA-seq we recovered a large number of differentially expressed genes (DEGs), although a significant portion of them had very small log2 fold changes. To focus our analysis on those that were likely to be most impacted we filtered our dataset to a minimum log2 fold change of 1 before further analysis.

Interestingly, we see a high level of variability in the DEGs affected between cell types, even given the similarity of fibroblast and MSCs (*17*) (Fig. 2B), with the majority of DEGs being unique to each cell type. This is also true even if we remove the minimum log2 fold change (Fig. S2). This suggests that these deletions likely have cell-type specific effects and may result in very different phenotypes depending on the cell’s specific context.

### Potential impact on mammoth phenotypes

We next wanted to see if we could infer some of the functions of these deletions, particularly as they may relate to arctic adaptations in woolly mammoths. To do this, we first screened our filtered DEGs to identify genes within pathways of interest (Fig. 2C), and then investigated what was known about perturbations in those genes in various cell and animal model analyses. We identified changes that broadly fall under three phenotypes: skin and hair adaptations; vascular adaptations; and metabolism and thermogenesis.

### Skin and Hair Adaptations

In comparison to their closest living relatives, Asian elephants, woolly mammoths have both denser hair coverage and multiple coats (layers of hair) to protect them from the arctic cold (*8*). We find DEGs which may be related to skin and hair adaptions within each deletion, although not in every cell type – MSC Deletion 3 (SOSTDC1 and BCL2); MSC Deletion 51 (FABP5); and Fibroblast Deletion 89 (IGFBP5). SOSTDC1 is involved in the regulation of the number, size and/or shape of hair, and is upregulated during the anagen phase in response to depilation-induced hair regeneration (*18*, *19*). BCL2 prevents apoptosis, and its upregulation has been shown to protect against hair cell death in the utricle (*20*). Likewise, we observe downregulation of IGFBP5 in our fibroblast Deletion 89 cells, which is a strong negative regulator of hair development and differentiation (*21*). Previous studies have also found that overexpression of IGFBP5 produces thinner and shorter hair with aberrant morphologies (*21*). The upregulation of SOSTDC1 of these in MSC Deletion 3, and downregulation of IGFBP5 in Fibroblast Deletion 89, suggest these deletions may contribute to the abundance and robustness of the mammoth’s coats.

We also identify FABP5 upregulated in MSC Deletion 51, which is involved in regulating water permeability. Knockouts of FABP5 result in much lower fatty acid content in the epidermis and impaired water-barrier function and keratinocyte migration (*22*). Likewise, this gene is expressed in sebaceous epithelial cells where it plays a role in sebaceous wax production (*23*). Sebaceous wax is believed to play a role in waterproofing and water loss, as well as thermoregulation (*8*, *24*). Mice which have been engineered to lack or have nonfunctional sebaceous glands suffer from abnormal hair growth/loss, skin lesions, and compromised thermoregulation (*24*). Notably, histological analyses of the few mammoth soft tissue remains that have preserved have revealed the presence of sebaceous cells, which are almost entirely absent in extant elephants (*8*, *25*).

### Vascular Adaptations

Vascular adaptations are also frequently observed in many arctic species. One of the best studied adaptations within this category is an increase in heart size, likely due to the need for more blood flow to maintain temperature and oxygen homeostasis (*26*, *27*). Although mammoth soft tissue has poor taphonomy, there are claims that mammoth hearts were larger than similar warm-adapted elephants of their age and size (*28*). Correspondingly, we observe genes involved in heart remodeling and angiogenesis in four experimental conditions – MSC Deletion 3 (IGF2BP2); MSC Deletion 51 (NRG1 and LAMA4); and Fibroblast Deletion 51 (HBEGF and PPP1R3A). Previous research has implicated IGF2BP2, NRG1, LAMA4, HBEGF, and PPP1R3A in heart morphology and metabolism, with the overexpression of many of these genes resulting in heart remodeling and hypertrophy (*29–33*). Additionally, IGFBP2 and PPP1R3A have also been shown to play a role in metabolic changes that accompany abnormal heart morphologies in mice (*31*, *32*).

Accompanying these changes, we also observe differential expression in genes related to angiogenesis – MSC Deletion 51 (LAMA4); and Fibroblast Deletion 89 (SEC14L3). SEC14L3, in conjunction with VEGF, has a role in promoting the migration of endothelial and vein progenitor cells and consequently angiogenesis, and its overexpression has been implicated in the proliferation and metastasis of tumors (*34*). Likewise, LAMA4 appears to be highly involved in blood vessel development. Mouse LAMA4 knockouts appear generally phenotypically normal, but exhibit frequent hemorrhages, and in cornea angiogenesis assays developed grossly distorted blood vessel architectures (*35*). Interestingly, while both genes are strong drivers of angiogenesis, we observe downregulation of SEC14L3 and upregulation of LAMA4 in their respective experimental conditions. Although the general trend here is still unclear, it’s possible these deletions are implicated in fine tuning angiogenesis in mammoths.

### Metabolism and Thermogenesis

The ability to sense and respond to cold temperatures via metabolic changes is probably amongst the most important cold adaptations. There is a suite of metabolic cascades that are triggered in response to cold nociception, and the ability to respond to cold stimuli via thermogenesis is highly dependent on the regulation of both glucose and lipid metabolic reserves (*36*). Not surprisingly, we recover many differentially expressed genes involved in adipocyte proliferation and lipid and glucose metabolism in four out of our six experimental conditions.

The bulk of our differentially expressed genes are involved in adipocyte differentiation and/or lipid metabolism – MSC Deletion 3 (IGF2BP2, PID1, and NPC1L1); MSC Deletion 51 (IGF2BP2, MSR1, DGAT2, DUSP26, and DDIT3); Fibroblast Deletion 51 (NR4A1, EGR2, and MAFB); and Fibroblast Deletion 89 (IRF4 and IGFBP5). We observe upregulation of IGF2BP2, PID1, DDIT3, and IRF4 in their respective experimental conditions, all of which except DDIT3 are positive regulators of adipocyte differentiation/proliferation (*37–40*). Likewise, we observe downregulation of NR4A1, EGR2, and MAFB all of which have previously been shown to act as negative regulators of adipocyte differentiation or size (*41–43*). Unsurprisingly, almost all these genes and additionally MSR1, DGAT2, NPC1L1, DUSP26, and IGFBP5 are also involved in lipid metabolism and controlling the rates of stored versus circulating lipids (*44–48*). For example, mice induced to overexpress DGAT2 do not appear to significantly different fat pads or body weights relative to controls but exhibit much higher levels of circulating plasma triglycerides (*47*). Likewise, MAFB deficient mice have higher body weights and amounts of body, due to large adipocytes and higher serum cholesterol levels (*43*).

Accordingly, many of the above genes have been shown to be implicated in the ability to sense and respond to cold in mouse models. GRIK2 (MSC Deletion 3) knockouts are unable to sense cold, but not cool, temperatures and have a sever reduction in cold-specific neurons (*49*). IGF2BP2 knockout mice have better thermoregulation than controls in the short term, but have fewer and smaller adipocytes and adipocyte precursors, and have diminished UCP1 levels, a key thermogenic protein (*38*). Interestingly, IRF4 knockouts have been shown to be more obese than wild type mice but exhibit increased cold intolerance likely due to the cold responsive nature of IRF4 (*37*).

### The Asian elephant genome contains 29 TP53 retrogenes

Although our RNA-seq results suggest that the deletions we engineered are likely involved in woolly mammoth cold adaptations, many of these phenotypes would only be measurable in organismal models. Towards this end we have primarily referenced literature on perturbations in these genes within mouse models, though we note that elephants are quite diverged from all common mammalian model species (*50*). To both validate our cellular models, and expand our understanding of elephant cancer resistance, we investigated the best-studied elephant genetic system – the expanded TP53 repertoire.

We searched the annotations of the Asian elephant reference genome and identified 29 retrogene copies of TP53 (RTGs). 27 of these RTGs are found on chromosome 27, while the remaining two are found on chromosomes 24 and 26. By comparison, the canonical copy of TP53 is located on chromosome 19. Most of the RTGs are approximately 1.1 kb in length, roughly the same length as the CDS of TP53 as would be expected of retrogenes, except for three (LOC126068247 – 64.1 kb; LOC126068267 – 32.5 kb; LOC126068248 – 4.9 kb; Fig. S3). When aligned and translated in the same frame as TP53, each RTG carries a premature stop codon, which primarily cluster in five main areas across the homology region (Fig. 3A).

**Figure 3.**
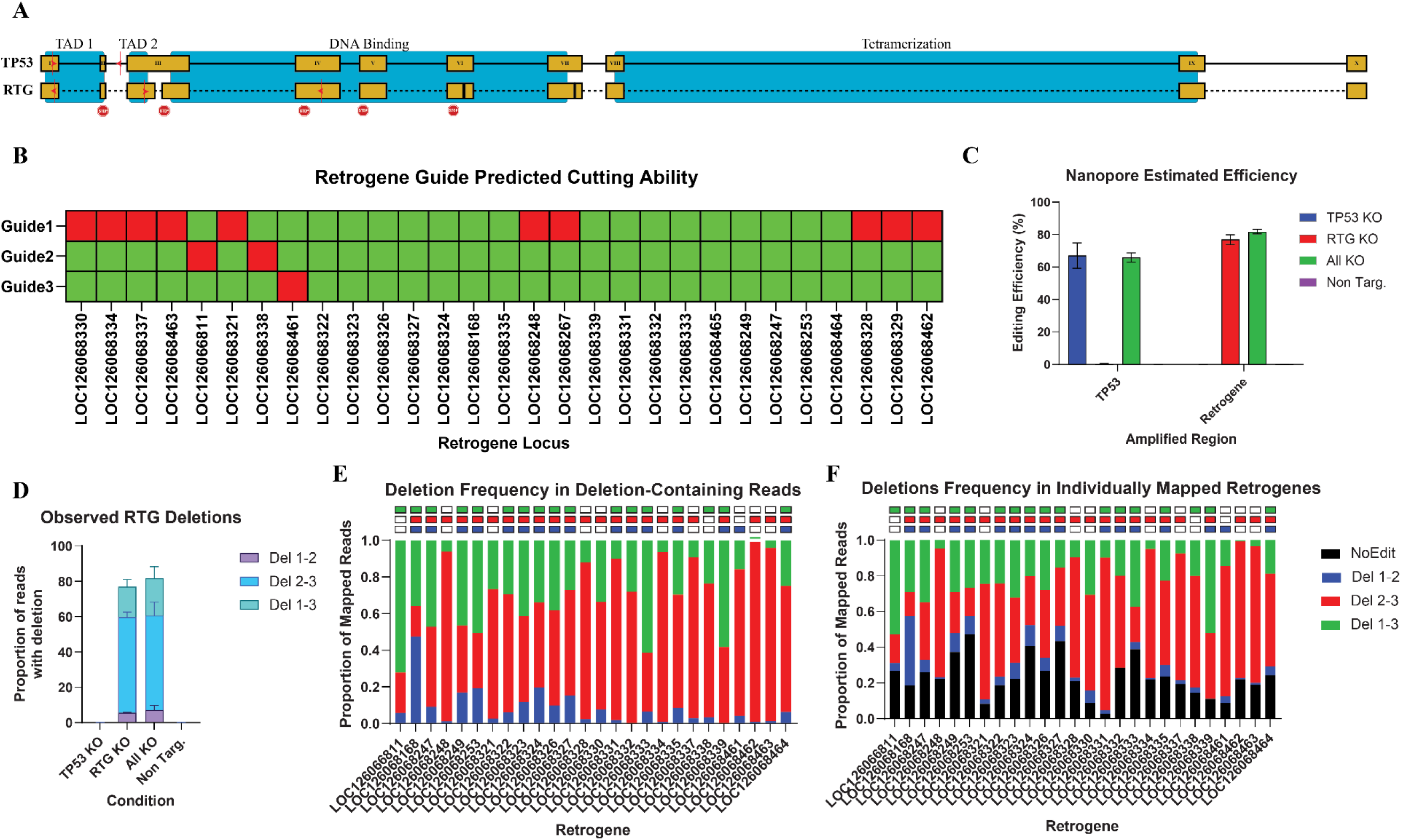
TP53 Editing. **(A)** Schematic showing an alignment of Asian elephant TP53 and a retrogene (RTG) consensus, with domains labelled based on homology to human TP53. Exons are shown in yellow, with introns indicated by black lines. Missing sequence is indicated with a dashed line in the RTGs. Additionally, we’ve included the positions of deletions greater than 3bp in the RTG consensus, and the approximate locations of stop codons observed across the 29 Asian elephant retrogenes. The locations and directions of guides are shown for both TP53 and the retrogenes. **(B)** Predicted guide activity against each of the 29 Asian elephant retrogenes, with green indicating the guide is expected to be functional against that target and red indicating non-functionality. **(C)** Observed editing efficiency of the expected deletions based on nanopore data. (**D)** The frequency of deletions observed in all reads mapped against the RTG consensus across different treatment conditions. **E.** The frequency of each deletion across the individual Asian elephant retrogenes as a proportion of all deletion-containing reads mapped to that locus, or all mapped read **(F)**. Amplicons which were identical across the non-edited regions were collapsed as they are bioinformatically undistinguishable. Error bars represent standard deviation in all plots.

### TP53 and RTG editing and validation

We attempted to design paired Cas9 guide RNAs (sgRNAs) to produce large, disruptive deletions in TP53 and all the retrogenes. We managed to identify two sgRNAs that specifically target TP53, which result in the partial loss of exon 1, the entirety of exon 2, and a premature stop codon after read through into intron 2 (Fig. 3A). We could not identify just two guides that were both predicted to have activity across all 29 retrogenes, so we designed a third guide to ensure each RTG was targetable by at least two sgRNAs (Fig. 3B; Fig. S4).

We electroporated fibroblast cells with Cas9 RNPs consisting of either the two TP53 guides, the three RTG guides, all five guides together (All), or a guide predicted to have no activity in the Asian elephant genome (Non Targ.). After approximately four days, we then collected cells, extracted DNA, and amplified a portion of TP53 and RTG genes flanking the expected editing sites. We observed high levels of editing across all three conditions when we examined the resulting amplicons on an agarose gel (Fig. S5). However, due to our PCR reaction amplifying all 29 loci simultaneously, we couldn’t assess which and how many of the RTGs were being edited.

To address this issue, we sent both PCR reactions for end-to-end nanopore sequencing. We quantified the number of observed deletions, using a custom script which marks the presence of a deletion if 90% of the bases within a defined region were absent. We note that this is likely an underestimate of the true editing efficiency, as this will only identify the perfect deletion product and miss any indels and/or incomplete loss of the flanked segment of DNA. We observed high editing efficiency in all conditions using either the TP53 guides (TP53 KO – 67%; All KO – 66%) or the RTG guides (RTG KO – 77%; All KO – 82%), and almost no editing in the Non Targ. condition (<0.1% at both loci) (Fig. 3C). We further examined the proportion of each deletion product observed in the RTG PCRs, and noted a clear pattern, wherein deletions between guides 2 and 3 were the most frequent, in line with expectations about the targeting scope of those two guides (Fig. 3D).

Next, we wanted to disentangle the editing on a per-locus level for the RTG data. We mapped the data to each unique RTG (collapsing those with identical sequences outside the editing region) and identified the proportion of each deletion at each locus. Although we generally observe a mix of all three deletions at each locus (some of which is likely due to cross mapping between the RTGs), we generally observe deletions in line with our expectations based on their targeting scope (Fig. 3E). Only two loci buck this trend, LOC126068338 and LOC126068461, both of which have a single point mutation at position 10 within the seed region of the guide (Fig. S4). Due to the importance of this region for guide specificity, we filtered these targets out; however, we note that at least at these two loci the guides appear to be permissive for editing. When combined with the per-locus reads that were not identified as containing a deletion (Fig. 3F), we observe an average of 76.8% editing efficiency across all unique loci.

### RNA-seq analysis reveals overlapping and unique TP53 and RTG responses

Given most literature on elephant RTGs has largely focused on how these retrogenes may interact with canonical TP53 and MDM2, we first decided to look at changes in expression of all direct downstream targets of TP53 (Fig. 4). In the absence of stress (i.e. 10µM Mitomycin C), we see relatively small changes in expression within most downstream targets with the exception of MDM2 and CDKN1A in both the TP53 and All KOs. Interestingly, we note that TP53 is not identified as significantly downregulated in the TP53 KO, suggesting maybe that nonsense-mediated decay is less pronounced in Asian elephants. In comparison, we observe very few differentially expressed genes in the RTG KO, and only TP53 that has a log2 fold change greater than 1. This suggests that the RTGs may play some role in either stabilizing or increasing the abundance of p53, although likely not directly by acting as a transcription factor as none of the RTGs have an intact DNA-binding domain (Fig. 3).

**Figure 4.**
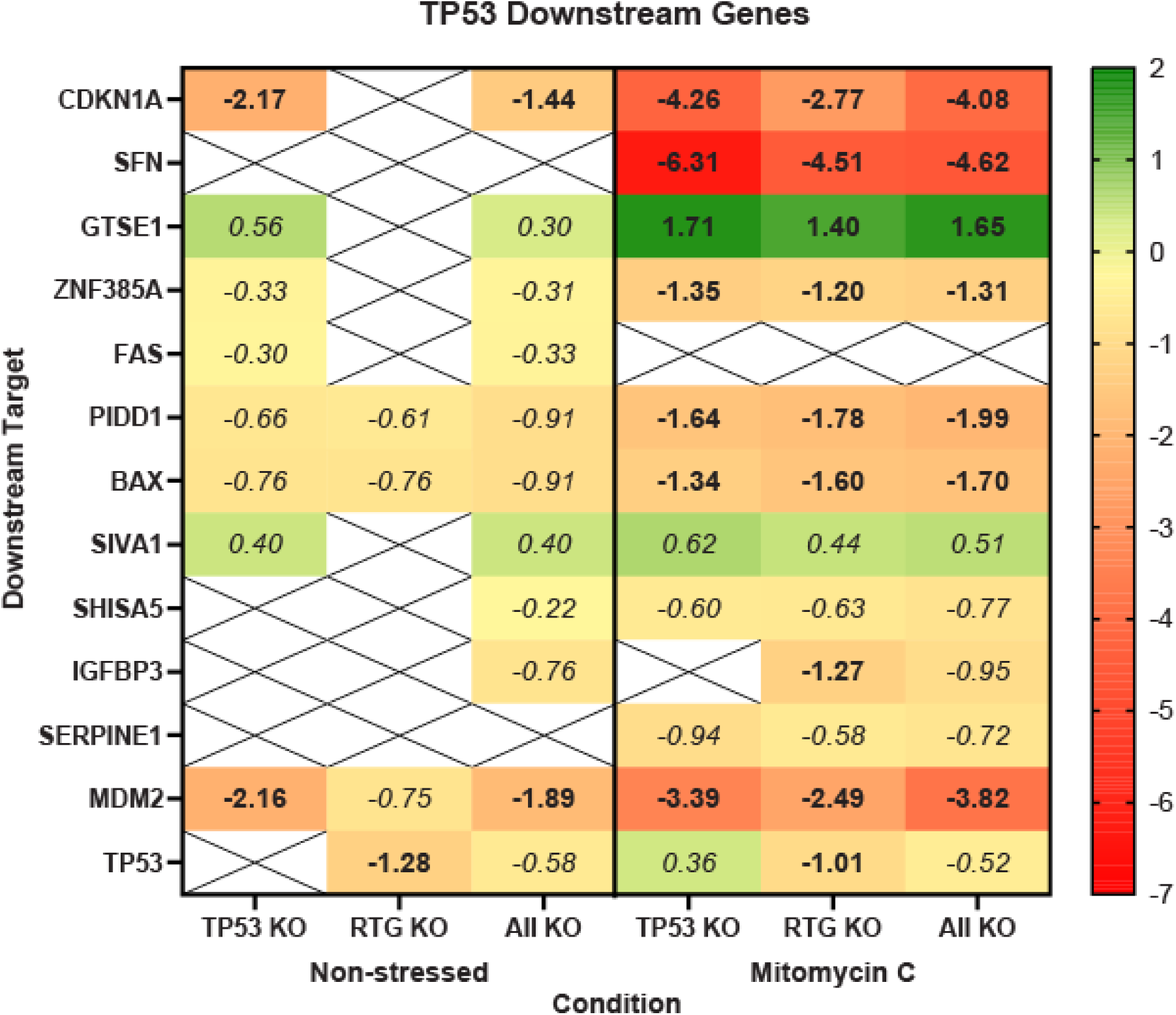
Expression of TP53 pathway genes. Log2 fold changes observed for genes immediately downstream of TP53 in the p53 signaling pathway (KEGG pathway hsa04115). Crosses indicate that no significant difference in expression was observed for that gene in a given condition. Genes in italics have an absolute log2 fold change of < 1 and were filtered out of enrichment analyses. Only downstream genes with a significant difference in at least one condition are shown.

In the presence of Mitomycin C, all three conditions have similar patterns in DEGs (Fig. 4), although typically smaller expression changes in the RTG KO. As before, we also observe that TP53 is significantly less expressed in the RTG KO and not in the TP53 KO. Additionally, IGFBP-3, a gene which can contribute to cell growth and apoptosis (*51*, *52*), is also downregulated in the RTG KO. We also observe significant downregulation of SFN across all three conditions in the MMC-treated cells, in line with this gene’s known tumor suppressor function of arresting cellular replication in response to DNA damage (*53*, *54*).

Next, we wanted to examine the different pathways enriched within each condition. Due to the large number of genes with small changes we filtered the dataset to a minimum absolute log2 fold change of at least 1. We then extracted the DEG lists for each condition and analyzed the overlap between them in either the non-stressed or MMC-treated cells (Fig. 5A). We performed BioPlanet pathway enrichment (*55*) on the 763 DEGs common to all three conditions treated with Mitomycin C. We observed significant enrichment in p53 signaling pathways, as would be expected of cells exposed to a DNA damaging agent, as well as enrichment in many DNA replication and cell cycle pathways (Fig. 5B).

**Figure 5.**
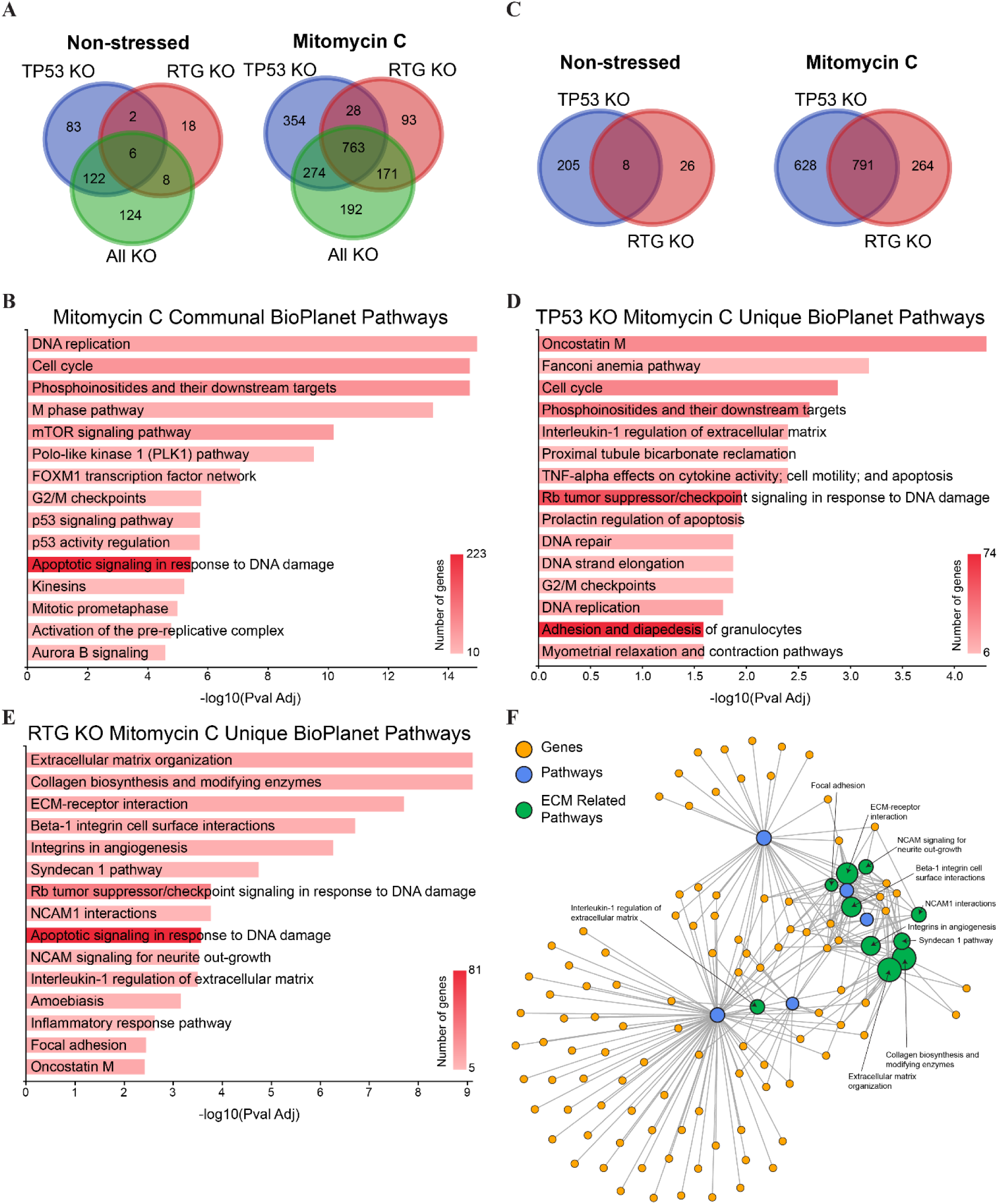
Pathway enrichment. **(A)** Differentially expressed gene (DEG) overlap observed between treatment conditions for cells in normal media or exposed to 10µM mitomycin C. Genes were filtered for a minimum absolute log2 fold change greater than 1. **(B)** BioPlanet pathway enrichment for the 763 DEGs common to all three conditions in the Mitomycin C treatment cells. **(C)** DEG overlap between TP53 KO and RTG KO cells in normal media or exposed to 10µM Mitomycin C. **(D)** BioPlanet pathway enrichment of unique DEGs in TP53 KO Mitomycin C treated cells. **(E)** BioPlanet pathway enrichment of unique DEGs in RTG KO Mitomycin C treated cells. **(F)** Pathway/gene cluster analysis for DEGs in the RTG KO Mitomycin C treated cells. ECM related pathways are colored green and labelled. Pathway clusters are scaled by their p-value.

We also examined the unique DEGs in the TP53 and RTG KO cells with Mitomycin C treatment (Fig. 5C). For the TP53 KO, we see enrichment in many pathways that would be expected given the known checkpoint and cell stress responses coordinated by TP53 (e.g. cell cycle and proliferation, DNA repair, etc; Fig. 4D) (*56–58*). By comparison, unique RTG KO DEGs are enriched in extracellular organization and signaling/interaction pathways (Fig. 5E), with many genes that contribute to multiple extracellular matrix related processes (Fig. 5F). This suggests that RTGs may play a unique protective role in regulating the tumor microenvironment and limiting metastasis (*59*). Additionally, the largest number of DEGs appear to belong to the apoptotic signaling in response to DNA damage pathway, a function that has previously been suggested for African elephant RTGs (*12*, *14*).

### Very few RTGs are likely to be expressed or functional

Given the unique function suggested by the RNA-seq of the RTG KO cells, we next wanted to see if we could investigate which RTGs might be responsible for the changes we observed. In none of the conditions, irrespective of treatment, did we detect differential expression of any of the RTGs.

Furthermore, we generally do not observe differences in the amount of RTGs expressed in response to Mitomycin C treatment (Fig. S10A), with the exception of the All KO. We do observe an increase in the amount of RTGs expressed in the RTG and All KO conditions, irrespective of Mitomycin C treatment (significance values not shown), although this may be a lingering transcriptional effect caused by double-strand break repair (*60*). However, we do note that all but three RTGs have very low read counts (Supplementary Table 6).

We next wanted to confirm that this pattern in RTG expression is not due to cell type or tissue specific. We download public RNA-seq datasets for Asian elephant ovary, salivary glands, lymph nodes, lung, thyroid, and lymphocytes, as well as examined some preliminary data generated with the same guides in epithelial and mesenchymal stem cells, and examined RTG expression patterns in these datasets. We observe the same general pattern, where the only three RTGs which show consistent expression are LOC126068247, LOC126068267, and LOC126068248 (Supplementary Table 7). However, these three RTGs are the three largest RTG genes (LOC126068247 – 64.1 kb; LOC126068267 – 32.5 kb; LOC126068248 – 4.9 kb) based on their annotations.

To examine what was driving the expression of these RTGs, we extracted RNA-seq mapped reads across each of these three loci and examined the frequency of reads across each locus. For two of these RTGs, we observe that most of the reads assigned to these RTGs map thousands of base pairs upstream of the region homologous to the TP53 CDS. In LOC126068247 and LOC126068248, we observe almost no mapped reads within the TP53 homology region either in our data or the publicly available datasets (Figs. S11-S14). By comparison, LOC126068267 also has peaks of high coverage upstream of the TP53 homology region but shows similar levels of coverage across the homology region in both datasets (Figs. S15-S16).

Given the lack of expression of most of the annotated retrogenes, we decided to compare those that might be functional in Asian elephants with African elephant retrogene 9 (RTG 9; KF715863), on which most previous studies have focused (*12–14*). We aligned the TP53 homologous regions of all Asian elephant RTGs, African elephant RTG 9, and the TP53 CDS, and built a maximum-likelihood phylogeny.

We observe that RTG 9 is most closely related to a clade containing two of our putatively expressed RTGs, LOC126068267 and LOC126068248 (Fig. S10B), further suggesting that these two RTGs might be responsible for the changes we observe. However, we do note that due to the high similarity of many of these RTGs, the bootstrap support values within our phylogeny are quite weak.

## Discussion

Proboscidean evolution can be characterized by a general increase in size from the 5 kg *Eritherium* to the 13-ton *Palaeoloxodon*, and a dispersal from the tropical wetlands of Africa into a wide variety of environments across the globe (*6*, *7*). Elephants offers a unique opportunity to examine these trends as they contain a highly evolved tumor suppressor system (*12*), and have closely related taxa adapted to two distinct environments – the arctic steppe tundra (woolly mammoths), and the tropical forests and grasslands of southeast Asia (Asian elephants) (*6*). Consequently, these taxa represent key candidates to understand both the adaptations necessary to expand into arctic environments and maintain cellular and genomic integrity with increased body size.

Studies in other taxa have suggested that a lot of adaptive evolution takes place on the level of gene regulation as opposed to protein function, possibly due to the reduced constraints on regulatory motif mutations (*2*, *5*). In this study we have investigated the regulatory role of some mammoth-specific deletions on arctic adaptations, and the expanded TP53 repertoire found in all elephants. We do note that all of these results should be viewed as hypotheses and testing their true impact would require the generation of animal models to fully examine their developmental and tissue-specific effects. However, as this is both currently not possible and ethical in elephants, we have generated many of these mutations within elephant cells to shed some light on their possible function.

We observe many differentially expressed genes that may be involved in arctic adaptations when we introduce our identified mammoth-specific deletions. Amongst these we see genes involved broadly with vascular system development, skin and hair, lipid metabolism, temperature sensing, and thermogenesis. However, we often fail to observe differential expression of genes containing or near our perturbations, although this result is likely affected by cell type (*61*). In fact, even in our larger mammoth deletions, we observe very little overlap in DEGs between MSCs and fibroblasts. Taken together, the results presented here are promising and provide some insight into mammoth adaptations but likely fail to fully capture the regulatory extent of these mutations, and which likely will not become evident until transgenic non-model species become commonplace or our ability to bioinformatically predict causative effects within a tissue-specific context drastically increases.

We also find that only three of the twenty-nine TP53 retrogenes are expressed across an extensive list of tissues and likely functional. Interestingly, two of the Asian elephant retrogenes most likely to be functional are closely related to African elephant RTG9, which has been shown to induce strong cellular responses to DNA damage (*12–14*) and apoptosis in mouse fibroblasts and human cancer cell lines (*12*, *14*). We also observe clear transcriptional differences between TP53 and RTG KO cells, including a significant downregulation of TP53 expression in our RTG KO cells, suggesting a novel function for elephant retrogenes. Additionally, we observe a differential enrichment of pathways when disrupting RTGs versus TP53, indicating that while there is a high amount of overlap between these genes, they also regulate many unique pathways. In particular, genes enriched within many extracellular organization and signaling pathways were observed in the RTG KO population. Some of these (e.g. extracellular matrix organization, angiogenesis) have been implicated as contributing to the establishment of tumor microenvironments, and are gaining increased importance as vital to cancer growth and spread (*59*). This may also explain the low prevalence of malignant cancers in Asian elephants (*62*).

Our work highlights the utility of cellular models to begin disentangling the complex effects of regulatory noncoding deletions and cancer resistance genetic networks in non-model species. However, in addition to the tissue-specific response limitations mentioned above, it is further hindered by our reliance on primary, non-immortalized cells, although this will likely be addressed as genetic resources for non-model species become more abundant (*63*, *64*). Of particular importance would be stable and readily available iPSCs, which would allow for differentiation of various tissues and more complex organoid models to better investigate the effects of both adaptive and cancer resistance mutations, especially where the generation of live animal models is not possible.

## Supporting information

Supplementary Tabbles

## Acknowledgments

We would like to thank the San Diego Frozen Zoo for sharing the fibroblast cell line. We would like to thank Zhengkuan Tang for discussions on RNA-seq experimental setup and analysis, and Ana Luiza Lemos Queiroz and Ramiro Perrota for assistance during early pilot experiments. We would also like to thank George Chao for guidance on nanopore analysis, and Sergiy Velychko for help during troubleshooting of the fibroblast cell culture techniques. Lastly, we would like to thank Roman Teo Oliynyk and Chun-Ting Wu, as well as the rest of the Church lab for helpful discussions throughout this project. **Funding**: EK and AAT were supported by a structured research agreement between Harvard Medical School and Colossal Biosciences. **Author contributions**: EK and GMC conceived the study with feedback from EM and AAT. EK and NB designed and conducted all wet-lab experiments with feedback and help from EM, AAT, and LL. EK and LL completed all bioinformatic analyses. AAT provided additional RNA-seq data for epithelial cell lines. EK wrote the first draft of the manuscript. All authors helped in the revision of subsequent drafts. **Competing interests**: For a complete list of G.M.C.’s financial interests, please visit http://arep.med.harvard.edu/gmc/tech.html. All other authors declare no other conflicts of interest. **Data and materials availability**: Bulk RNA-seq data for all mammoth deletions as well as TP53 conditions with and without mitomycin was deposited on NCBI (BioProject Accessions: PRJNA1147676 and PRJNA1217465). Bioinformatic scripts and tools created for this work are available at: https://github.com/EmilKarpinski/RTG_TP53_MammothDel_Analysis.

## Supplementary Materials

Materials and Methods

Supplementary Tables 1 to 7

Figs. S1 to S16

Table S1

References (65-85)

## Methods

### Elephant Genomic Dataset

All bioinformatics analyses for deletion identification used previously published datasets. We used datasets for eleven mammoths (*Mammuthus*), thirteen Asian elephants (*Elephas*), and twenty-two African elephants (*Loxodonta*) (Supplementary Table 1). Published datasets were aligned to the *Elephas maximus* reference genome (GCF_024166365.1) using BWA aln (*67*).

### Mammoth Deletion Identification

A summary of the mammoth deletion pipeline is available in Fig. S1. Briefly, the *Elephas maximus* reference genome was split into individual contigs. These were then fed into GenerateWindows.awk (WinSize = 25; WinSlide = 10; AmbProp = 0.2), to generate windows across the genome and filter out any with more than 20% missing data (i.e. N’s). We then generated read depth files for each mammoth/elephant alignment using samtools depth (-a to output all positions) (*74*), and split these by contig. Windows were then cross referenced with depth files using another CoverageCalc.awk to extract windows containing no mapped reads (StdevMultiplier=99), and adjacent windows concatenated using ConcatonateWindows.awk. Windows with zero coverage identified in *Loxodonta* and *Elephas* were then combined using bedops (*75*), then were used to filter the identified mammoth deletions using bedtools intersect (-v flag to retain windows which are uniquely in the mammoth dataset) (*76*).

We also attempted to predict transcription factor binding sites (TFBS) upstream of elephant genes. We first extracted the transcription start side and direction of each protein coding gene in the *Elephas maximus* reference genome, then generated intervals for 1kb upstream of each gene. These intervals were extracted using the samtools faidx program (*74*), and TFBS predicted using TFBStools (*77*) in custom Rscript (TFBSTools_JASPAR2014.R). Identified TFBS were then cross referenced against filtered mammoth deletions using bedtools intersect (*76*) to identify putative cis-regulatory deletions present in woolly mammoths.

During the course of this project, a separate ancient DNA deletion pipeline was published (*15*). We also performed deletion prediction using the published pipeline using the same datasets above. We examined the overlap between our two datasets using bedtools intersect (*76*).

### TP53 identification and primer design

We extracted and aligned the canonical TP53 locus as well as all 29 retrogenes (RTGs) from the *Elephas maximus* reference genome (GCF_024166365.1). For the RTGs, we trimmed the alignment to include only the ∼1.1kb segment overlapping the exons in TP53. We then designed two primers to specifically amplify the TP53 gene (Supplementary Table 2). For the RTGs, we designed seven forward and four reverse primer variations, all amplifying the same locus, but containing all possible binding site mutations to minimize amplification bias (Supplementary Table 2). These primers were made up to 10µM stock solutions and equal volumes of all forward and all reverse primers combined to produce master forward and reverse stock solutions.

For all other deletions, we generated at least four primers flanking the region containing the deletion ensuring similar melting temperatures across target sites, and that all primers were outside of the region targeted by Cas9. For each locus, we tested all possible combinations of forward and reverse primers and chose those that produced the cleanest and most reproducible amplifications (Supplementary Table 2).

### Guide Design

We generated all possible guide RNAs (gRNAs) within TP53 from the start of first exon to the end of the last exon using the generate_kmery.py script distributed as part of GuideScan2 (*78*). All guides were then screened for predicted off-targets against the entire *E. maximus* reference genome using GuideScan2 with a maximum mismatch distance of 3 (-m 3), and with the option to output all guides (-t-1). The output was then further filtered using a custom python script (Cas9_MMFiltering_V1_1.py) and scoring matrix to filter potential gRNAs based on how many mismatches they possessed relative to potential off-target sites and where along the spacer those mismatches occur. From the remaining guides we selected two guides (Supplementary Table 1) that cut within exon 1 and between exon 2 and 3 respectively to generate TP53 knockouts as these disrupt the TAD1 domain and should interfere with proper splicing resulting in a premature stop codon.

To design guides for the RTG knockouts, we generated all possible guides against the trimmed RTG alignment for all 29 retrogenes using the generate_kmery.py script as before. We then screened all guides for those that were predicted to have activity to the most possible targets and identified two gRNAs that were each predicted to have activity against all but three retrogenes, which were only targetable by either one of the two guides (Supplementary Table 2; Fig. 3). We then subsequently chose one additional guide that targeted those three remaining retrogenes to ensure that each retrogene had at least two gRNAs.

We also identified three genes that had consistent expression across our MSC and epithelial cell line: FN1, EIF3I, and CD109. For each of these we designed all possible guides as above and filtered for off-targets against the *E. maximus* reference genome using GuideScan2 and filtered as above. We then selected four guides for each spaced roughly equally apart across a 1kb upstream region of each gene.

For all other deletions we extracted the *E. maximus* reference sequence ±25bp from the start or end point of each deletion using samtools faidx (*74*) and generated all possible guides within those regions using generate_kmery.py. We then filtered the automatically generated guides for off-targets using GuideScan2 and the python script as above. In the case multiple guides were possible for each deletion we chose one that would result in a double-stranded break closest to the original endpoint of each deletion.

### Cell culturing and media

Fibroblast cells were grown in a custom media formulation with the final concentrations: 50% FGM-2 (Lonza FBM Basal Media (CC-3131) supplemented with FGM-2 SingleQuots supplements (CC-4126)); 20% heat-inactivated fetal bovine serum (ThermoFisher, A5670801); and 30% Mem Alpha (Gibco, 12571063). Mesenchymal stem cells were grown in a custom media formulation with final concentrations: 79% DMEM High Glucose + GlutaMax (ThermoFisher 10566016); 20% heat-inactivated fetal bovine serum (ThermoFisher, A5670801); and 1% MEM Non-Essential Amino Acids Solution (ThermoFisher 11140050). Epithelial cells were grown in a custom media formulation with final concentrations: EGM-2 (Lonza Basal Medium (CC-3156) supplemented with EGMTM-2 SingleQuotsTM Supplements (CC-4176)); 40% DMEM High Glucose + GlutaMax (ThermoFisher 10566016); and 10% heat-inactivated fetal bovine serum (ThermoFisher, A5670801). Cells were grown in an incubator at 37°C with 5% CO_2_.

### Cas9 RNP Electroporation

All gRNAs were ordered from Synthego and resuspended to 100µM in 1X TE buffer. Cas9 nuclease was ordered from IDT (catalog # 10007807). All nucleofections were conducted using the Lonza P4 Primary Cell 96-well Kit (catalog # V4SP-4096) using codes CA-137 (fibroblasts) or EN-150 (epithelial cells), or the P3 Primary Cell 96-well Kit (catalog # V4SP-3096) using code CA-137 (MSCs).

For the TFBS, synthetic, and large-mammoth deletions, approximately 100-200k cells were used for each 20µl transfection reaction, with a 3µM final concentration of each guide (i.e. both upstream and downstream of the deletion) and 1µM of Cas9 nuclease per guide (i.e. a 3:1 ratio of gRNA:Cas9). RNPs were pre-complexed in nucleofection solution for 20 minutes at room temperature then electroporated into cells using the above codes.

We conducted pilot studies for the seven identified TFBS deletions upstream of genes that showed consistent expression in our MSC line, although only one guide was used for deletions upstream of LOC126074853 and LOC126068891 due to the small size of these deletions. We also conducted pilot studies using all three synthetic deletion genes (FN1, CD109, EIF3I), as well as for the six smallest intragenic and seven smallest intergenic large mammoth-specific deletions. Deletions were carried out alongside negative control containing just Cas9 and/or a non-targeting gRNA RNP which has no significant homology to the *E. maximus* genome. All experimental conditions for pilot work were conducted in duplicate and in MSCs. We subsequently chose a subset of these deletions which generated a high number of DEGs for our final analysis. These deletions were done as above, except with four replicates each and accompanied by 3-4 negative controls using a non-targeting gRNA. For Deletions 3, 51, and 89 we also attempted to introduce these deletions into fibroblast and epithelial cells, however the epithelial cells failed to recover from the nucleofections and were ultimately discarded.

TP53 deletions were done as above, except with 400-500k cells per transfection. Cells were transfected with either both TP53 gRNAs, all three RTG gRNAs, both TP53 and RTG gRNAs (“All” condition), or a non-targeting gRNA (Supplementary Table 1). For the All nucleofection, gRNAs were first combined and then concentrated using a Savant SpeedVac DNA 120 Concentrator (Thermo Scientific), before being resuspended in premade nucleofector and supplement solution. Additionally, due to the presence of possible TP53 mutations that accumulate with prolonging passaging, a second separate nucleofection was conducted as above prior to the Mitomycin C stressed RNA-seq. In both cases, all treatment conditions were conducted using three biological replicates. Due to their greater robustness, their ease of culture and use, and the availability of younger passage cells, we elected to conduct these experiments only in our fibroblast line.

Following nucleofection cells were immediately transferred into 6-well plates containing prewarmed media and allowed to recover for 4 days. Media was changed 24 hours post nucleofection and every 2-3 days thereafter.

### Editing Efficiency

During the first passage (approximately 4 days post nucleofection), about 10% of the cells were collected and DNA extracted using the DNeasy Blood and Tissue Kit (Qiagen) using the manufacturer’s recommended protocol. For the TP53-editing extractions we amplified both the TP53 locus spanning the expected knockout deletion, and the retrogene loci. For all other extractions we amplified the locus spanning the targeted deletion and examined these loci across the negative controls as well. We preformed 25µL PCR reactions with final concentrations of 1X Phusion Hot Start Flex 2X Master Mix and 500nM of each primer, using between 1-3µL of template. Cycling conditions were: 98°C for 3 minutes; 12 cycles of 98°C for 15 seconds, then 69°C for 60 seconds; 25 cycles of 98°C for 15 seconds, 67°C for 30 seconds, then 72°C for 45 seconds; followed by a final extension at 72°C for 5 minutes. Resulting PCR amplicons were purified over MinElute PCR Purification columns (Qiagen) using the recommended protocol and visualized on E-gel EX 2% Agarose gels (Invitrogen) to confirm the presence or absence of any large deletions. We examined the relative fluorescence of our products using the Image Lab 6.1 (Bio-Rad). We then calculated a minimum editing efficiency (the relative fluorescence of our expected deleted product, relative to everything else in the lane) and a maximum editing efficiency (the relative fluorescence of all non-wild-type bands within each lane).

To confirm efficiency estimates and examine RTG deletion distribution in our TP53-editing experiments, we sent each of our TP53 and RTG amplicon pools for nanopore sequencing using the Premium PCR service at Primordium to get full length sequences of our DNA fragments. For each sample, raw reads were first filtered using a custom python script (FastqFilter.py) to between the maximum and minimum expected sizes +/- 10% respectively. Mapping was carried out using BBMap (*66*) using default parameters against either the TP53 amplicon, an RTG consensus amplicon, or a multi-fasta with 26 RTGs (RTG amplicons with identical sequences outside of the editing region collapsed).

To calculate editing efficiency, we used a custom python script (CigarDel.py) which uses a region file containing the deletion regions of interest, a minimum deletion proportion (i.e. the proportion of bases mapping to the deleted region that need to be missing), and the CIGAR string of each read to evaluate the presence or absence of various deletions in the data. For TP53 we analyzed the presence of the expected deletion if Cas9 induced a double-strand break at both sgRNA targeting sites. For the RTG we examined the presence of any of the three possible deletions between two guides. All analyses were carried out with a minimum deletion proportion of 0.9.

To examine the distribution of deletions at each locus we identified the names of reads containing deletions and extracted those that mapped to each locus. We then cross-referenced this with the deletions identified against the RTG consensus to obtain read distributions for reads mapped to all RTGs or the collapsed RTG loci.

### Bulk RNA-seq

Approximately 300-500k cells were collected, flash frozen on dry ice, and shipped to Genewiz for bulk RNA sequencing within ten days of transfection (TP53) or 2-3 weeks (all other deletions).

Expression of transcripts was done using Salmon (v1.8.0 (*69*)) using a custom index consisting of all mRNAs and non-coding RNAs annotated within the *E. maximus* reference genome (GCF_024166365.1), with all 29 TP53 retrogene sequences added. The salmon index was generated using the keepDuplicates flag to avoid collapsing any RTGs. Salmon quant files were analyzed using DESeq2 (*79*) as part of a custom Rscript to analyze samples in a pairwise fashion (i.e. one set of biological replicates against the controls at a time). Differentially expressed genes (DEGs) were identified with an adjusted p value of less than 0.05 (padj < 0.05). No minimum log2 fold change was applied for initial analysis of TP53 pathway genes. A minimum absolute log2 fold change of 1 was used for all other analyses. Venn diagrams and DEG overlap was generated using a web based custom venn diagram tool (https://bioinformatics.psb.ugent.be/webtools/Venn/), and pathway enrichment preformed using GeneCodis4 (*80*).

### Retrogene expression in public datasets

To ensure the expression patterns we observed were not tissue/cell type specific, we examined the expression of our retrogenes in some preliminary shared data from mesenchymal stem cells and epithelial cells. These cells contained deletions generated using the same guides as in this study, but the epithelial cells were also subject to various small molecule treatments. We also downloaded public RNA-seq datasets for Asian elephant ovary (SRX15554240), salivary glands (SRX15554239), lymph nodes (SRX15554238), thyroid (SRX15554237), lung (SRX15554236), and lymphocytes (SRX1423033), and analyzed it as above.

### Depth plots

To examine where reads from each of the three RTGs showing repeated expression were originating, we mapped the data for our datasets and the public datasets back to the Asian elephant reference genome using STAR (*81*). We then extracted read depths along the full annotated length of RTGs LOC126068247, LOC126068267, and LOC126068248 using samtools depth (*82*). For our dataset we averaged expression values within biological replicates and plotted the data in Microsoft Excel.

### Maximum likelihood phylogeny

We download African elephant retrogene 9 (KF715863.1) to investigate a possible relationship between this retrogenes and the Asian elephant retrogenes. We then aligned the retrogene to the Asian elephant TP53 CDS and truncated the RTG 9 sequence to contain only the homologous sequence to TP53. We then aligned the truncated RTG 9 sequence, Asian elephant TP53, and all 29 Asian elephant retrogenes using muscle (v5.1) (*83*). We generated a maximum likelihood phylogeny using IQ-Tree (*84*), using ModelFinder to identify the best supported substitution model (*85*), and 100 bootstraps (-b 100).

