## Supplementary Information for "Engineering elephant models of cold adaptation and cancer resistance"

**Supplementary Figure 1. Mammoth Deletion Pipeline Overview.** High level overview of the steps involved in identifying deletions in the proboscidean datasets, transcription factor binding site prediction, and their overlap.

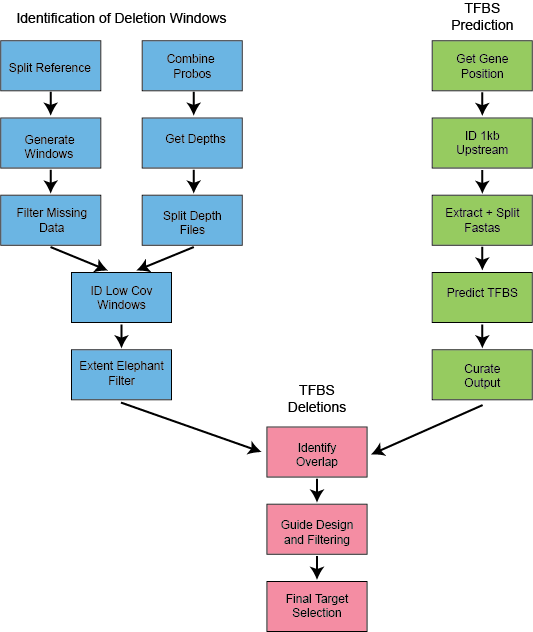

**Supplementary Figure 2. DEG Overlap.** Overlap of identified DEGs between MSC and Fibroblasts with the same deletion with no log2 fold filtering.

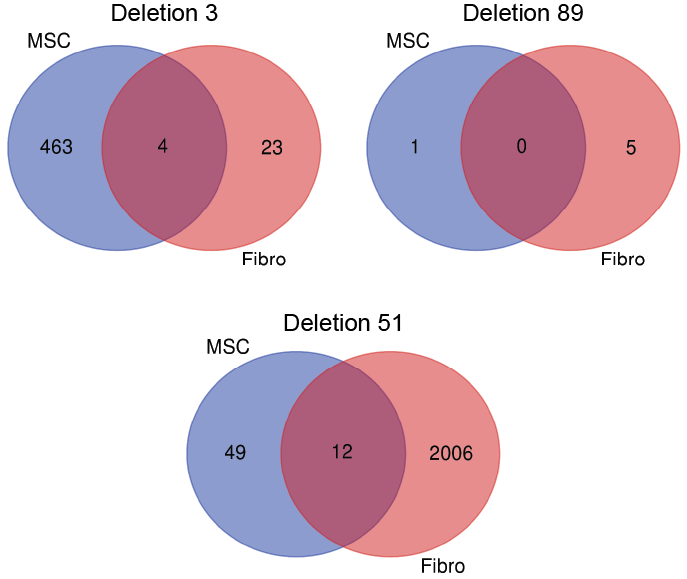

**Supplementary Figure 3. RTG characterization. A.** Length distribution of Asian elephant RTGs and the TP53 CDS. **B.** DNA or protein identity for the RTGs relative to TP53. **C.** DNA identity of the RTGs to the TP53 and to each other.

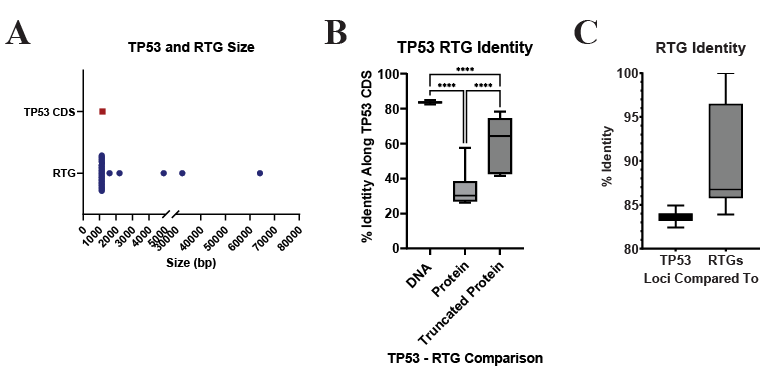

**Supplementary Figure 4. RTG guide alignments.** Alignments of each Asian elephant RTG with the three RTG sgRNAs. Red shading indicates guides predicted to be outside the targeting scope of that guide.

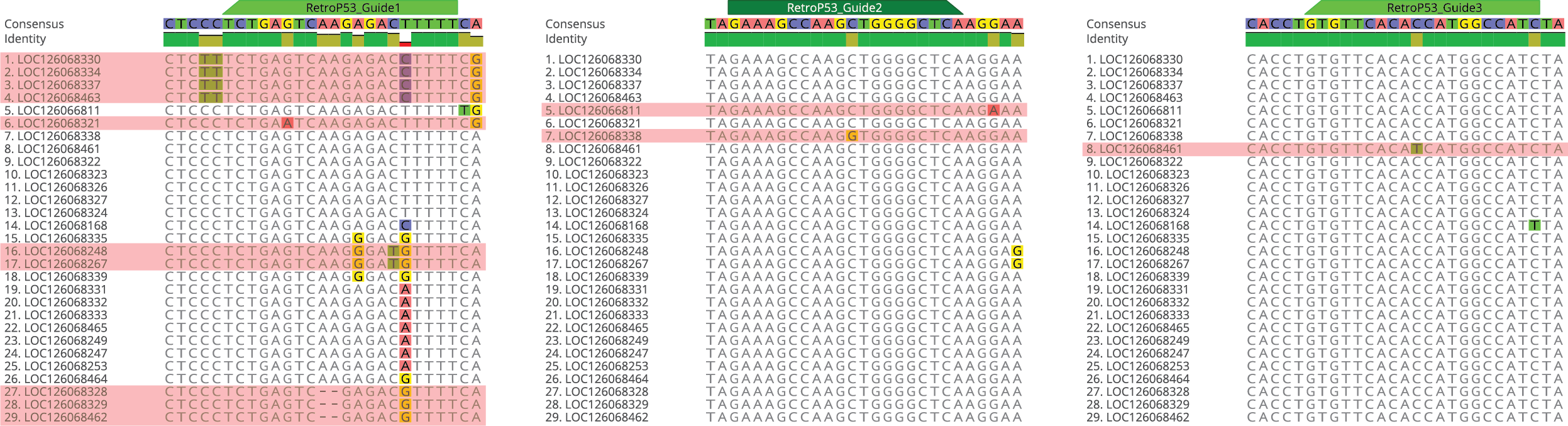

**Supplementary Figure 5. Gel electrophoresis editing efficiency. A.** Sample gel showing quantification of TP53 and/or RTG editing in one biological replicate for each of the four conditions. **B.** Editing efficiency following our initial nucleofection and following the redone nucleofection prior to Mitomycin C treatment and RNA-seq (to account for possible TP53 mutations). Editing efficiency was calculated both as the relative fluorescence of the expected deletion band relative to all other bands (Minimum), or as the relative fluorescence of all non-wild-type bands (Maximum). Errors bars represent standard deviation.

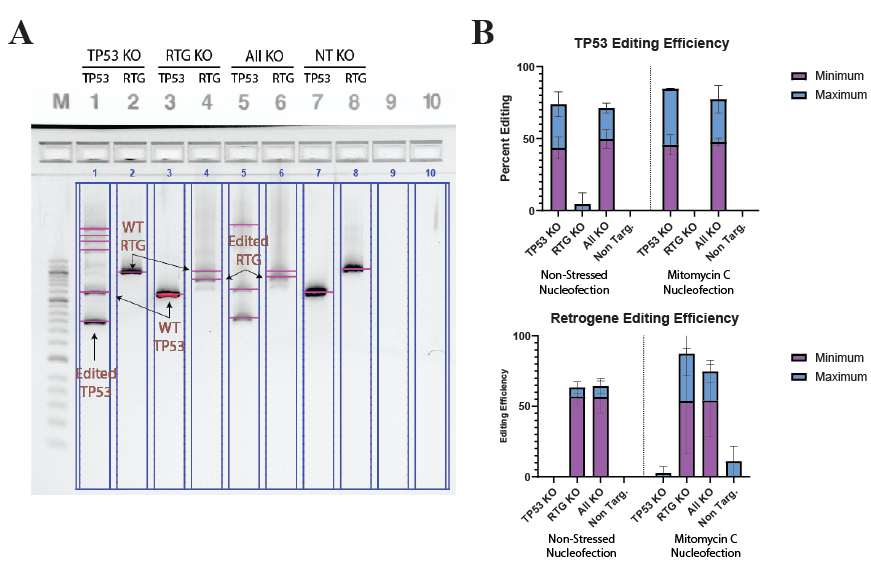

**TP53 Growth, Viability, and RT-qPCR**

As a preliminary test of the TP53/RTG deletion functionality, we examined the growth rate, viability, and the expression of key TP53 pathway genes using RT-qPCR. Approximately four days following transfection, we seeded 50k cells from each condition into four 24-well plates and observed the growth and viability of the cells over approximately 4.5 days. We observed no significant difference in the growth or viability (Fig. S6) of the cells across this time. However, we do note that the TP53 KO wells appeared to reach confluency a day before the other conditions suggesting faster growth, which is consistent with our experiences handling these cells in subsequent passages.

Next, we wanted to see if we could observe a functional effect of the knockouts we generated. We cultured 50k cells in the presence of increasing concentrations of Mitomycin C (MMC), a DNA damaging agent (Lee et al., 2006; Sulak et al., 2016), and examined the expression of p21 (CDKN1A) and MDM2, two direct downstream targets of p53, and an upstream regulator, ATR (Fig. S6). We observed significant decreases in expression of MDM2 and p21 at all non-zero Mitomycin C concentrations across each condition, and almost no significant differences in ATR. However, we were surprised to see that the response in the RTG KO was so similar to that of the TP53 and All KOs. We extracted DNA from the same line of cells one passage later and PCR amplified the TP53 and RTG loci again. We observed a drop in Sanger sequence quality across the TP53 amplicons in some of the biological replicates (Fig. S7), generally consistent with the location of the RTG guides. This suggests there may be some off-target editing in the RTG samples that becomes enriched during routine passaging of the RTG KO cells, particularly if these TP53 mutants also possess a potential growth advantage as would be suggested by our growth curve.

To minimize the effect of any off-target mutations on our RNA-seq results, we sent cells soon after nucleofection (within ~10 days). Despite this we observe the presence of off-target edits at the homologous location of the RTG guides within the TP53 transcripts (Fig. S8). Interestingly, we see significant enrichment of these TP53 mutations in our MMC-treated cells. This suggests that these TP53 mutations might confer a growth advantage to the cells as was also suggested by our growth curve analysis.

**Supplementary Figure 6. Cell characterization and expression. A.** Growth rates of cells with TP53, RTG, or combined knockouts, or treated with a non-targeting guide. **B.** Cell viability observed during the growth curve for each condition. **C.** The expression of two downstream (p21 and MDM2) and one upstream (ATR) genes within the TP53 pathway. Expression levels for the RTG KO are shown with a dashed line due to the possible presence of TP53 off-target mutations. Significance levels: p ≤ 0.05 (*); p ≤ 0.01 (**); p ≤ 0.001 (***); p ≤ 0.0001 (****). Errors bars indicate standard error of the mean in all plots.

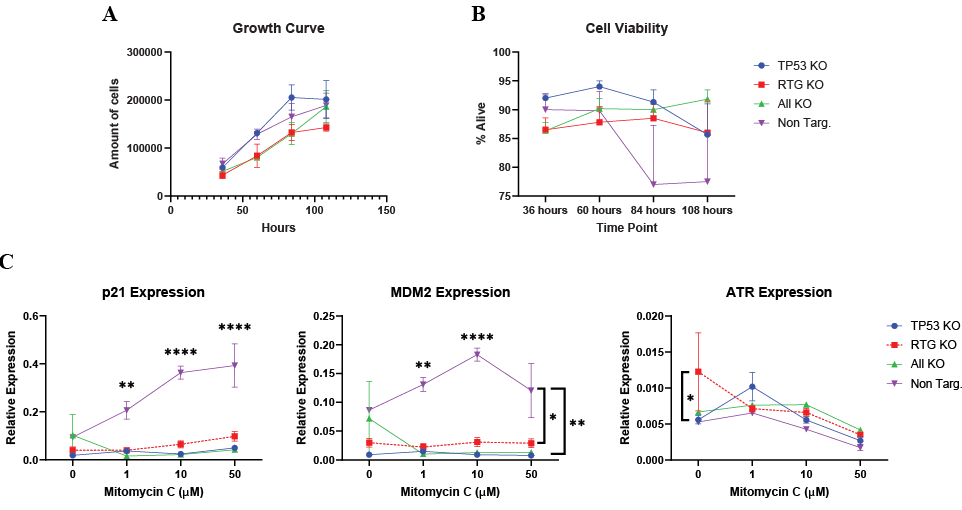

**Supplementary Figure 7. Potential TP53 off-target mutations.** Sanger sequencing read for the TP53 locus in two biological replicates of the RTG KO one passage following RT-qPCR. RTG guide are shown in red, and aligned to their homologous regions in TP53. Traces are zoomed out so the quality of the entire read (blue) can be seen. The trace in RTG KO – 2, shows a drop in quality roughly consistent with the position of RTG guide 1, suggesting possible off-target editing.

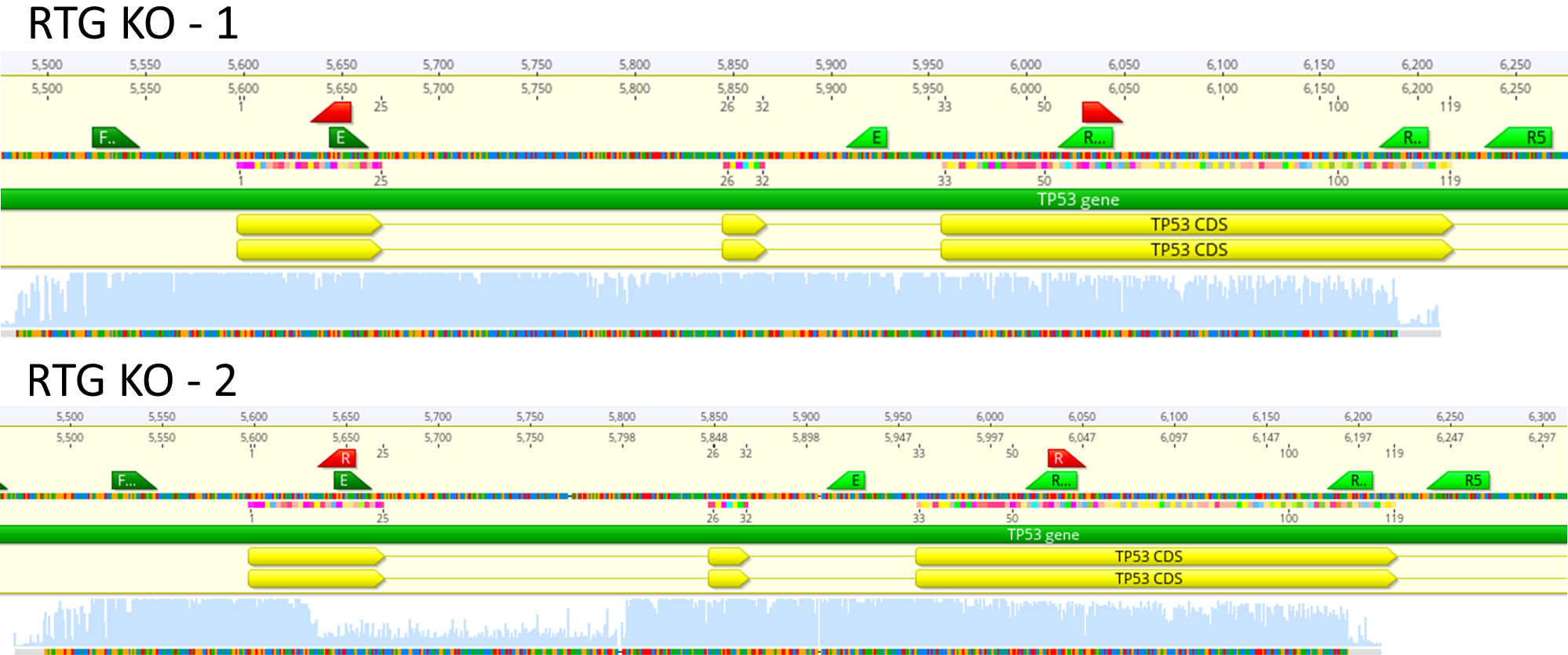

**Supplementary Figure 8. Cross-target estimation.** Estimation of cross activity within TP53 with the RTG-targeting sgRNAs estimated from the RNA-seq data mapped to the Asian elephant genome. The proportion of reads containing indels causing a frameshift is shown for the homologous region in TP53 for retrogene guide 1 (**A**), guide 2 (**B**), and guide 3 (**C**). All errors bars represent standard deviation.

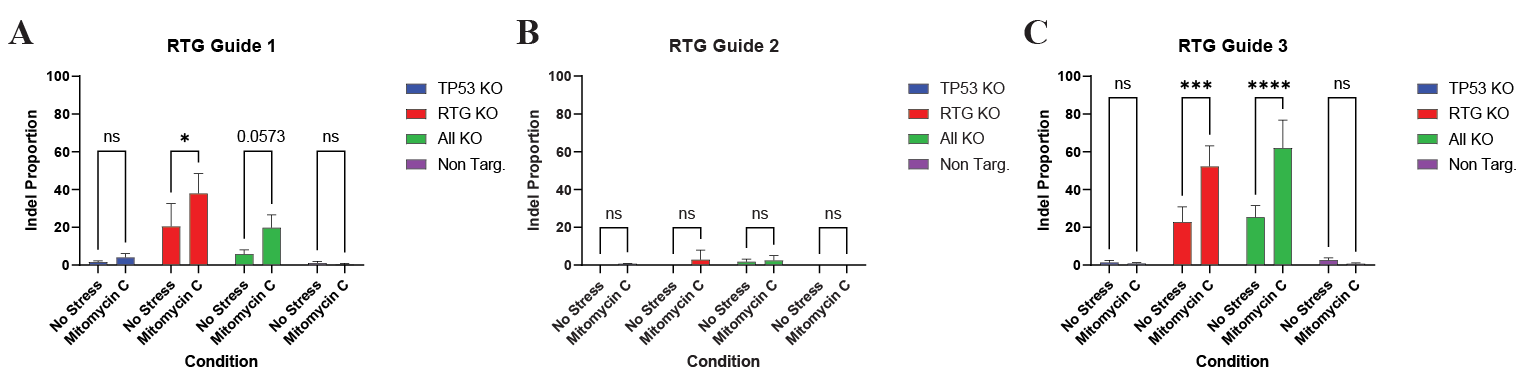

**TP53 RTG Synthetic Dataset**

To examine whether we could differentiate reads originating from TP53 and the retrogenes, and that our RNA-seq results were not due to mismapping between the RTGs and the canonical TP53 gene, we generated a synthetic dataset, consisting of 1,000 2x150 bp reads from TP53 and each of the RTG loci. We then used this dataset to compare the amount of reads assigned to each locus when mapped back to the genome using BBMap (Bushnell, 2022), BWA-MEM (Li and Durbin, 2009), or STAR (Dobin et al., 2013), followed by quantification with Salmon (Patro et al., 2017), or via direct quantification by Salmon. We find that all four methods gave similar results, with BBMap performing the worst, possibly due to how it assigns multi-mapped reads to the first locus in the reference (Fig. S9). In light of these results, we conclude that the decrease in TP53 expression is likely not a bioinformatics artifact due to poor mapping.

**Supplementary Figure 9. Read assignment during our synthetic mapping test. A.** The number of unique loci the reads from each RTG were assigned to. The ideal expectations, one assigned reference, is shown by the dashed red line. **B.** The proportion of LOC126068249 reads which mapped to different loci. **C.** Read counts as assigned by Salmon either from data mapped to the Asian elephant reference genome followed by Salmon quantification, or quantified directly by Salmon.

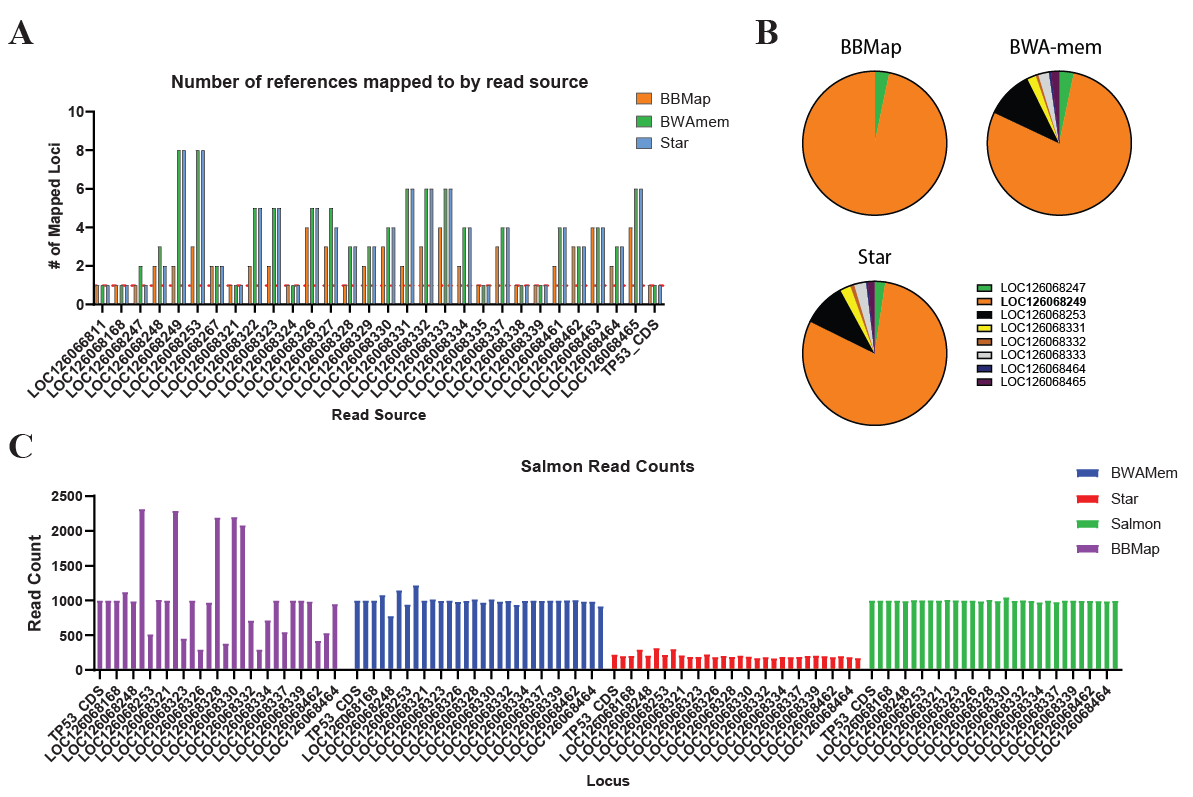

**Supplementary Figure 10. RTG expression. A.** Expression of RTGs in each treatment condition in cells exposed to 10µM Mitomycin C or not. Errors bars indicate standard deviation. **B**. Maximum likelihood phylogeny showing the relationship between Asian elephant TP53 and the RTGs, as well as African elephant RTG9. Three tree was rooted using Asian elephant TP53. Green circles indicate nodes with bootstrap support ≥ 90%, while numbers above nodes represent bootstrap support for select nodes. The three Asian elephant RTGs which are most commonly expressed, and African elephant RTG9 are indicated in orange and blue respectively.

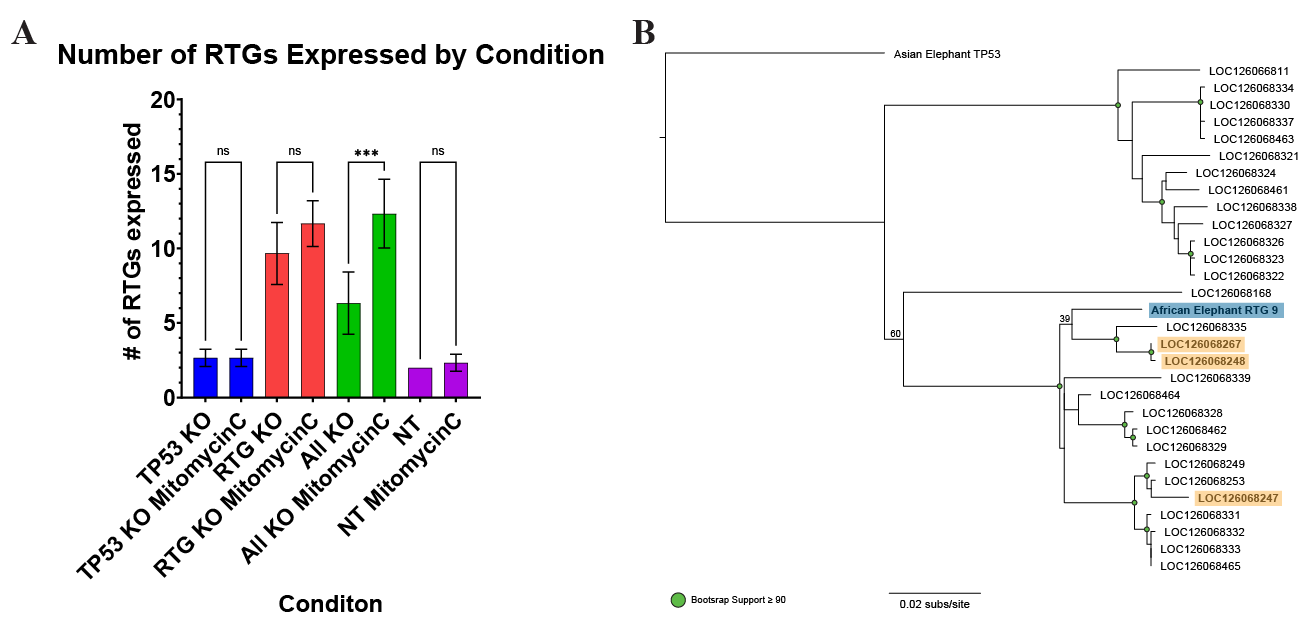

**Supplementary Figure 11. Read depth along LOC126068247**. **A**. Read depth along the entire annotated sequence of LOC126068247 using data in our datasets. **B**. A zoomed in version of the full plot showing only the TP53 homology region.

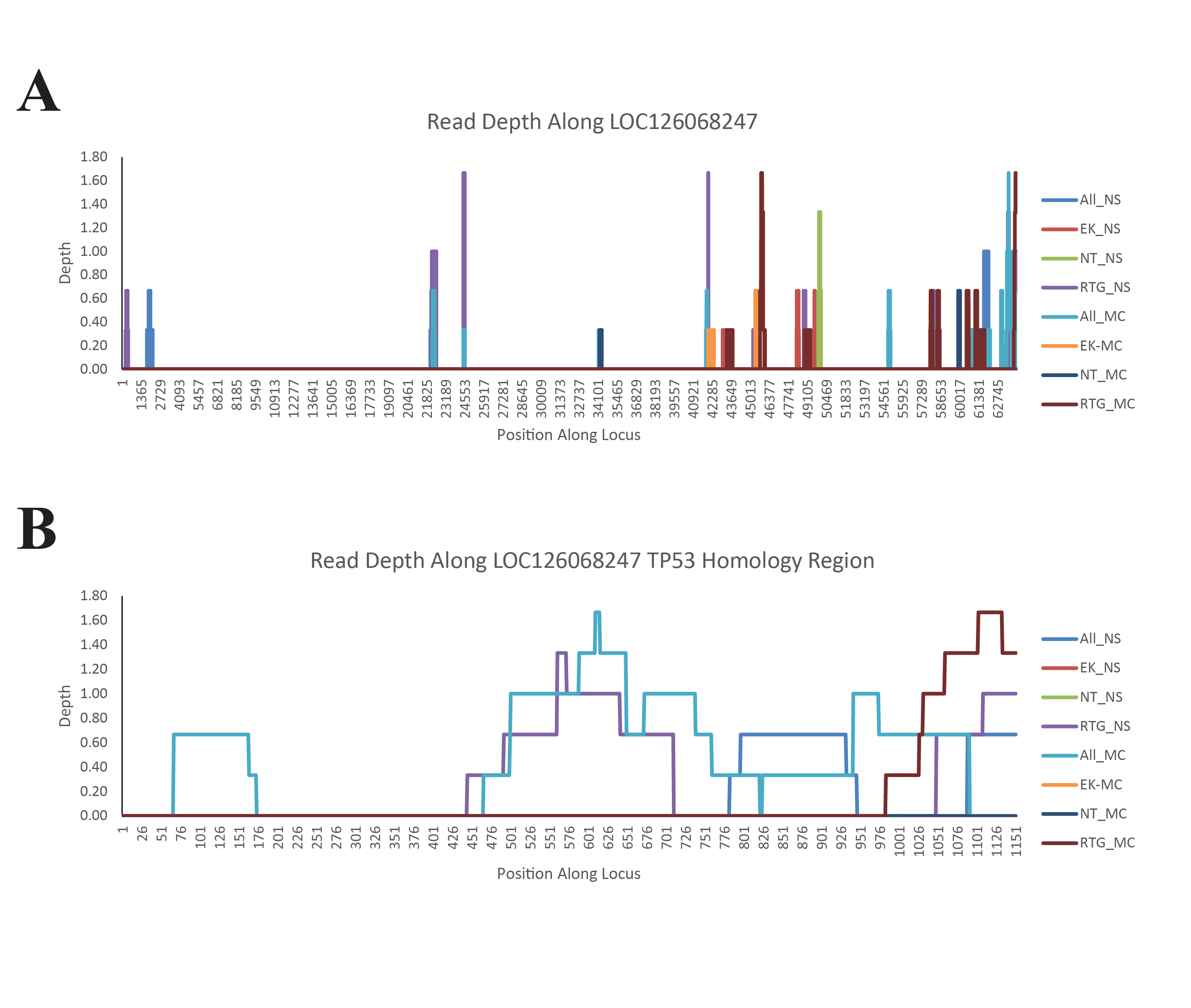

**Supplementary Figure 12. Read depth along LOC126068247**. **A**. Read depth along the entire annotated sequence of LOC126068247 using publicly available datasets. **B**. A zoomed in version of the full plot showing only the TP53 homology region.

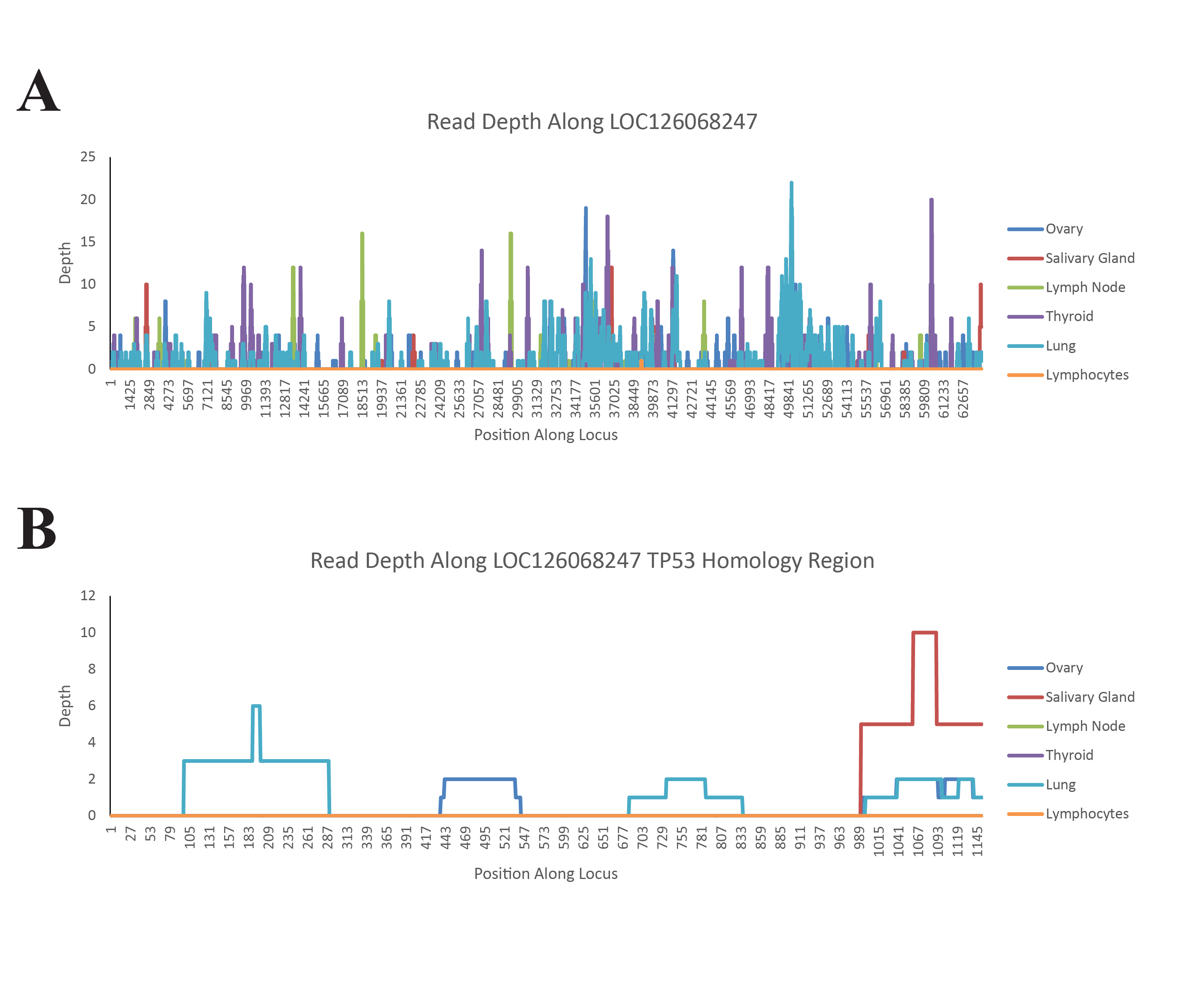

**Supplementary Figure 13. Read depth along LOC126068248**. **A**. Read depth along the entire annotated sequence of LOC126068248 using data in our datasets. **B**. A zoomed in version of the full plot showing only the TP53 homology region.

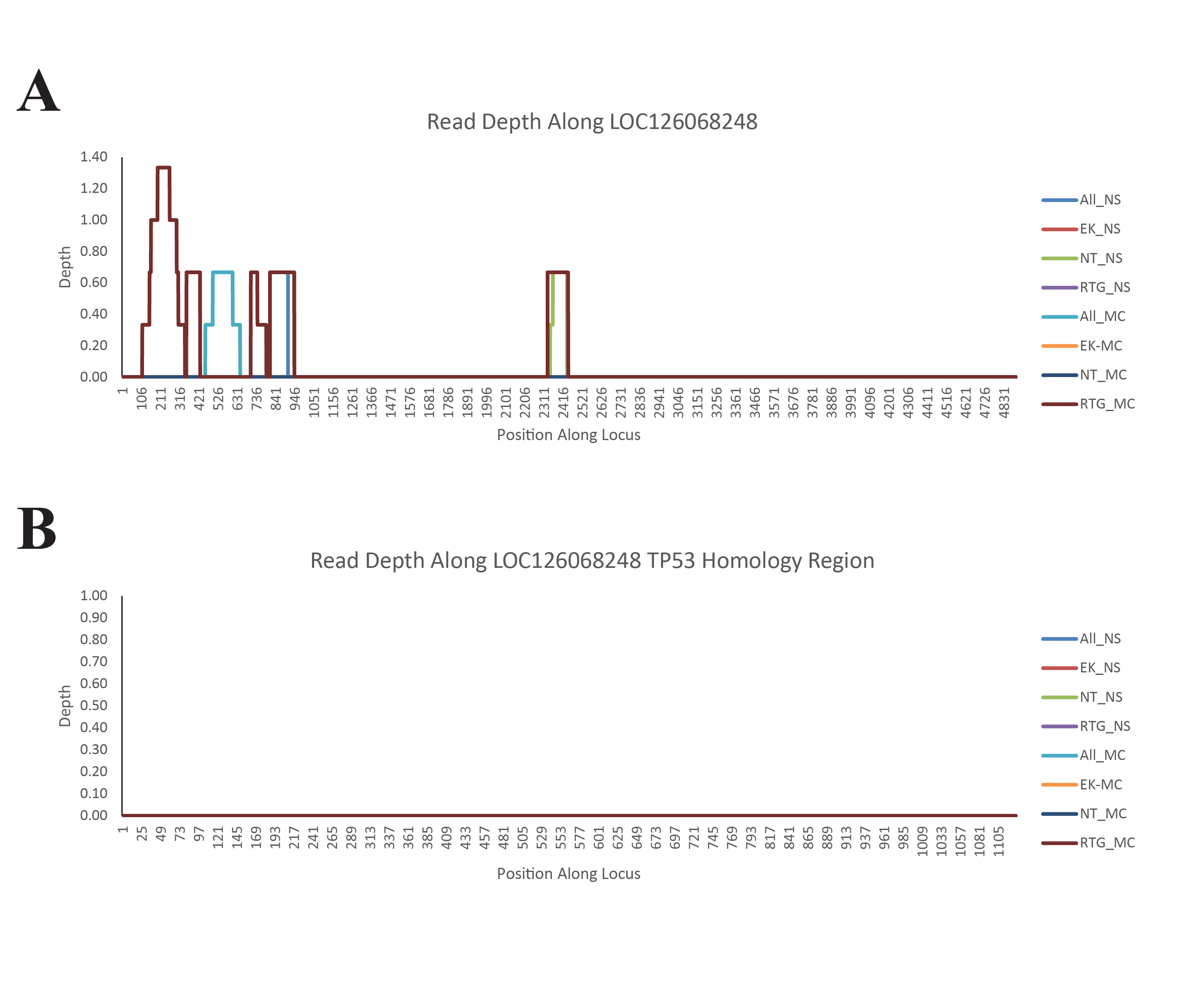

**Supplementary Figure 14. Read depth along LOC126068248**. **A**. Read depth along the entire annotated sequence of LOC126068248 using publicly available datasets. **B**. A zoomed in version of the full plot showing only the TP53 homology region.

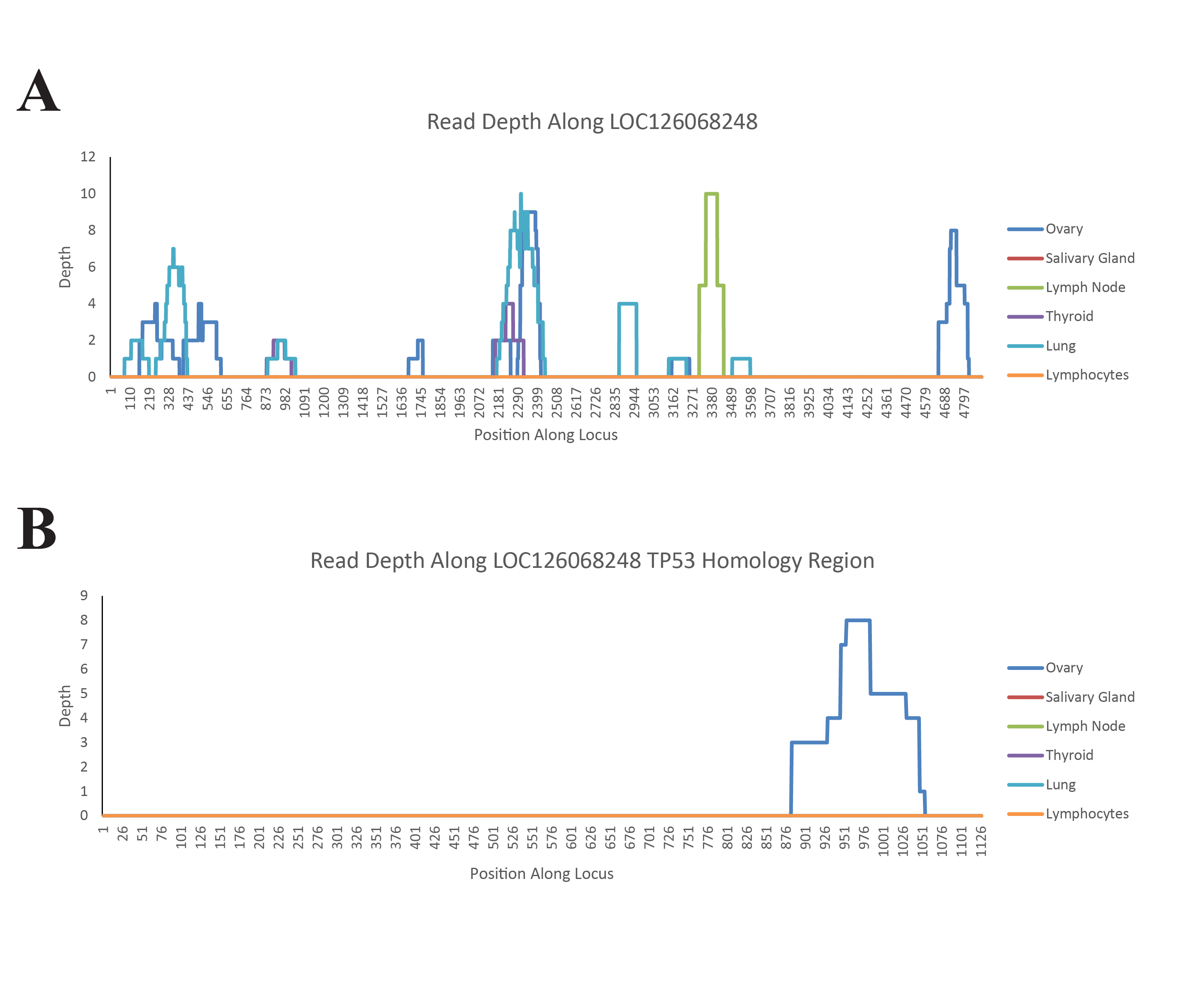

**Supplementary Figure 15. Read depth along LOC126068267**. **A**. Read depth along the entire annotated sequence of LOC126068267 using data in our datasets. **B**. A zoomed in version of the full plot showing only the TP53 homology region.

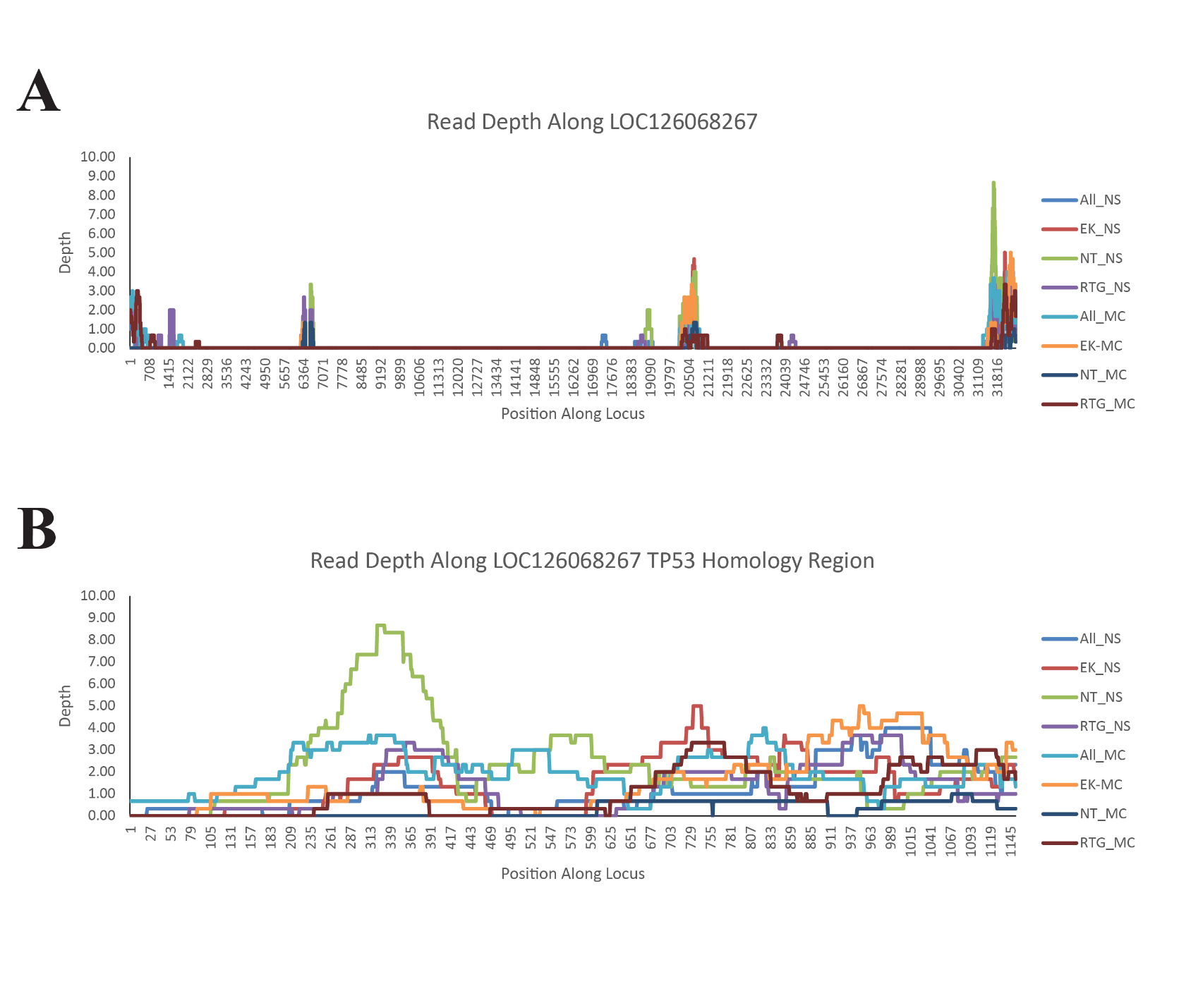

**Supplementary Figure 16. Read depth along LOC126068267**. **A**. Read depth along the entire annotated sequence of LOC126068267 using publicly available datasets. **B**. A zoomed in version of the full plot showing only the TP53 homology region.

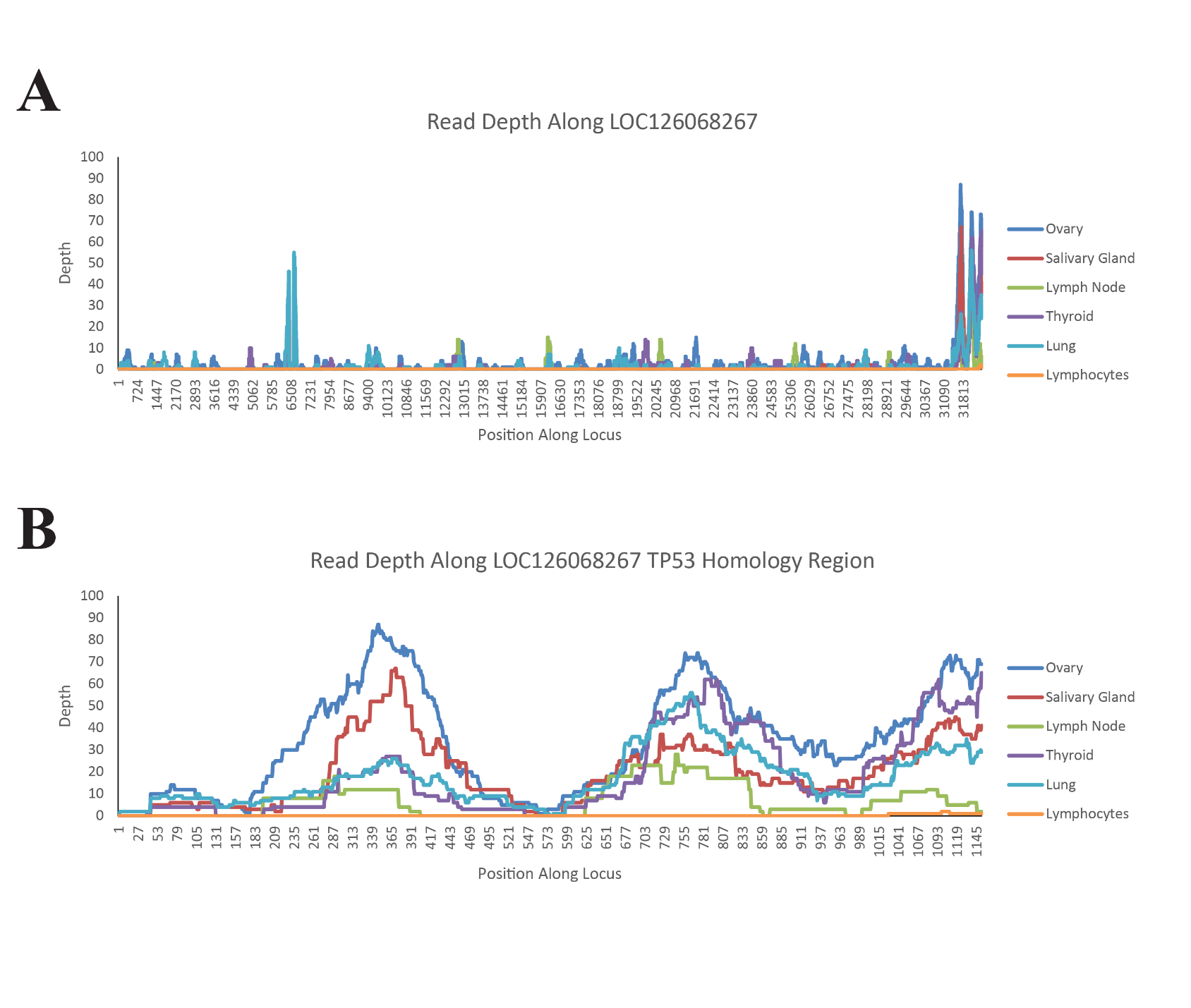

**Clonal Expansion**

Although compounded by a relative lack of studies and optimization relative to mouse and human cell lines, elephant cells have proven difficult to immortalize and grow (Fukuda et al., 2016; Pearson et al., 2021). We originally attempted to produce clonal lines with wild-type cells from our MSC line, as well as MSCs edited to contain either Del 3, Del 51, Del 89, or treated with a non-targeting guide using established protocols (Housden et al., 2015). After 20 days we observed no colonies in any well, and even our two wells seeded with 100 cells each (to identify the right plane during microscopy) failed to grow during this time. However, we reasoned that since TP53 mutations often precede cancer development, and many cancers naturally immortalize, we may be able to exploit this feature in our knockouts to produce isogenic lines.

To examine this, we treated cells with either 0µg/mL, 0.5µg/mL, or 1µg/mL EMS for 16 hours (Munroe and Schimenti, 2009), and then performed eight passages, seeding 10% of the detached cells, to outgrow fast growing and potentially immortalized cells. All 1µg/mL cells died within the first few passages, as did all the TP53 KO conditions and the Non Targ. 0µg/mL EMS treatment. After eight passages, the outgrown cells were collected and 1,000 from each condition were seeded into a 10cm^2^ dish, and colonies picked over 1-3 weeks. We observed between 4 and 26 well-defined circular colonies in the RTG and All KO conditions treated with 0µg/mL or 0.5µg/mL, in comparison to only one in the Non Targ. 0.5µg/mL condition (Table S1). Of these we picked three colonies each from the All 0 µg/mL, All 0.5µg/mL, and RTG 0 µg/mL plates, two colonies from the RTG 0.5µg/mL, and the only colony from the Non Targ. 0.5µg/mL plate. We attempted to expand colonies, but unfortunately most of the cells in most of these lines appeared quite sick and exhibited very slow growth. One of the RTG 0.5µg/mL colonies managed to propagate an additional four passages within the 10cm^2^ dishes, but slowed down immensely thereafter and became senescent.

**Table S1. Clonal Expansion.** The number of well defined circular single-cell colonies observed after 1-3 weeks of growth. Fields with a dash, indicate conditions that did not survive to colony propagation.

|  | EMS Concentration (µM) | | |
| --- | --- | --- | --- |
|  | 0 | 0.5 | 1 |
| TP53 KO | - | - | - |
| RTG KO | 4 | 18 | - |
| All KO | 26 | 16 | - |
| Non Targ | - | 1 | - |
